# Epigenetic state determines inflammatory sensing in neuroblastoma

**DOI:** 10.1101/2021.01.27.428523

**Authors:** Adam J. Wolpaw, Liron D. Grossmann, May M. Dong, Jessica L. Dessau, Patricia A. Brafford, Darya Volgina, Alba Rodriguez-Garcia, Yasin Uzun, Daniel J. Powell, Kai Tan, Michael D. Hogarty, John M. Maris, Chi V. Dang

## Abstract

Immunotherapy has revolutionized cancer treatment, but many cancers are not impacted by currently available immunotherapeutic strategies. Here, we investigated inflammatory signaling pathways in neuroblastoma, a classically “cold” pediatric cancer. By testing the functional response of a panel of 20 diverse neuroblastoma cell lines to three different inflammatory stimuli, we found that all cell lines have intact interferon signaling and all but one lack functional cGAS-STING signaling. However, toll-like receptor (TLR) signaling, particularly through TLR3, was heterogeneous. Six cell lines showed robust response, five of which are in the mesenchymal epigenetic state, while all 14 unresponsive cell lines are in the adrenergic state. Genetically switching the adrenergic BE2(c) cell line towards the mesenchymal state fully restored TLR responsiveness. In responsive cells, TLR3 activation results in the secretion of pro-inflammatory cytokines, enrichment of inflammatory transcriptomic signatures, and increased tumor killing by T-cells *in vitro*. Using single cell RNA sequencing data, we show that human neuroblastoma cells with stronger mesenchymal signatures have a higher basal inflammatory state, demonstrating intra-tumoral heterogeneity in inflammatory signaling that has significant implications for immunotherapeutic strategies in this aggressive childhood cancer.

## Introduction

In order to be effective, immunotherapies require cytotoxic immune cells to traffic to and enter the tumor microenvironment, recognize tumor cells, and kill them. These requirements provide a number of potential mechanisms for intrinsic or acquired immune evasion. For instance, with immune checkpoint blockade (ICB), resistant tumors tend to have lower pre-existing infiltration with cytotoxic T cells^1^ (trafficking and entry) and lower mutation rates^2, 3, 4^ (tumor recognition). The level of tumor cell inflammatory signaling has the potential to affect trafficking and entry of immune cells through the secretion of cytokines and tumor recognition through antigen presentation. Indeed, lower baseline inflammatory signaling has also been associated with ICB resistance^4, 5^ and response can be restored in some tumor models by activating tumor cell inflammatory signaling^6, 7^. Increasing inflammatory signaling has the potential to enhance other immunotherapies as well, including adoptive cellular immunotherapies and antibody-based therapeutics that require antibody-dependent cellular cytotoxicity (ADCC) for function.

Understanding which inflammatory sensing pathways are intact in a given tumor is an important initial step in determining how to activate cellular inflammatory signaling. These sensing pathways include the recognition of interferon, which can be secreted from both immune and non-immune cells, as well as the detection of molecular patterns typically found in microbial infection. Such patterns are recognized by a large group of receptors, including toll-like receptors (TLRs), often collectively referred to as pattern recognition receptors. These include proteins that detect dsRNA (TLR3, MDA5, RIG-I), ssRNA breakdown products (TLR7 and 8), CpG-rich (TLR9) or cytosolic DNA (cGAS), and lipopolysaccharides (TLR4), among others^8^. In tumors, products capable of stimulating such sensors can be released by dead and dying cells^9^, produced by activation of endogenous retroviral elements in response to therapy ^10^, exposed by breakdown of the nuclear envelope^11^, or transferred through exosomes from tumor or stromal cells^12, 13^. Some of these receptors have been shown to be critical to an immunologic response to anti-cancer therapy, including TLR3 with chemotherapy^14^ and cGAS with radiation therapy^11, 15^. Exogenous agonists of these pathways are being incorporated into clinical trials, particularly in combination with ICB. Thus, knowledge about the responsiveness of these pathways in cancer may contribute to our understanding of the basal inflammatory state of the tumor, suggest potential avenues to alter that state, and identify biomarkers for which agonist might be more effective in a given tumor.

Neuroblastoma accounts for ∼15% of childhood cancer mortality in addition to significant morbidity among survivors^16, 17^. Improved therapies are needed, particularly for high-risk disease, including the subset with amplification of *MYCN* (v-myc avian myelocytomatosis viral oncogene neuroblastoma-derived homolog)^17, 18^. Neuroblastoma is a classically “cold” tumor with a lack of T-cell infiltration, very low MHC I expression, and a low tumor mutation burden^19, 20, 21, 22^, all of which are even more pronounced in *MYCN*-amplified disease^23, 24, 25, 26^. Published studies to date have shown little to no efficacy with ICB^27, 28^. CAR T-cell therapies have demonstrated limited efficacy, thought to be in part due to poor penetration or function in the tumor microenvironment^29^. However, an antibody against the disialoganglioside GD2 is an immunotherapy that is FDA-approved for use in neuroblastoma despite significant off-tumor toxicity due to GD2 expression on pain nociceptors. This therapy improves survival when used in combination with cytokines in patients in first remission^30^ and has a ∼40% response rate when used in combination with cytokines and chemotherapy in relapsed disease^31, 32^. This demonstrates that the immune system can play a role in treating neuroblastoma, but expanding the benefit of immunotherapy to a greater percentage of neuroblastoma patients remains an important challenge.

One recent advance in the understanding of neuroblastoma biology is the discovery that neuroblastoma cells can exist in two different epigenetic states defined by super enhancer elements enriched in one state or the other^33, 34, 35^. There is a less-differentiated state described as either mesenchymal or neural crest cell-like and a more differentiated state described as adrenergic or sympathetic noradrenergic (herein referred to as MES and ADRN). These states are thought to co-exist within human tumors and can spontaneously interconvert, though cell lines tend to stably adopt one state or the other, with some exceptions. In human primary neuroblastomas, the ADRN state comprises the majority of the tumor at the time of diagnosis^33,34^. Evidence from human tumors^33, 36^ and mouse models^37^ indicates that the MES state may be enriched in relapse and in metastatic disease^38^, and MES cell lines are more chemo-resistant *in vitro*. Hence, the ability to specifically target the MES subpopulation may prove important in both treating and preventing relapsed disease. To our knowledge, connections have not been made between ADRN and MES states and inflammatory signaling or response to immunotherapy.

To identify ways to increase inflammatory signaling in neuroblastoma, here we define the responsiveness of a large panel of neuroblastoma cell lines to different inflammatory stimuli, test how those differences impact tumor-immune interactions, and identify determinants of response, revealing immune vulnerabilities of neuroblastoma.

## Results

### Neuroblastoma lacks basal inflammatory signaling

We first asked if the immunologically “cold”^19, 20, 21^ nature of neuroblastoma correlated with a lack of tumor cell inflammatory signaling. We compared expression of the Hallmark Inflammatory Response signature^39^ across a large number of tumor cell lines using RNA-seq data from the Cancer Cell Line Encyclopedia project^40^. Analysis of 991 cell lines across 37 tumor types showed that neuroblastoma has the lowest expression of this signature as well as several other inflammatory signaling signatures (Figure 1a and S1a), demonstrating that neuroblastoma cells have strongly suppressed inflammatory signaling. We then asked how *MYCN* amplification status impacts this signature. Similar to published reports^23^, we found that inflammatory signaling is further decreased in the *MYCN-*amplified subset of both cell lines and tumors compared to those without *MYCN* amplification, which corresponds to less infiltration with cytotoxic T-cells in tumors (Figure 1b,c).

**Figure 1:**
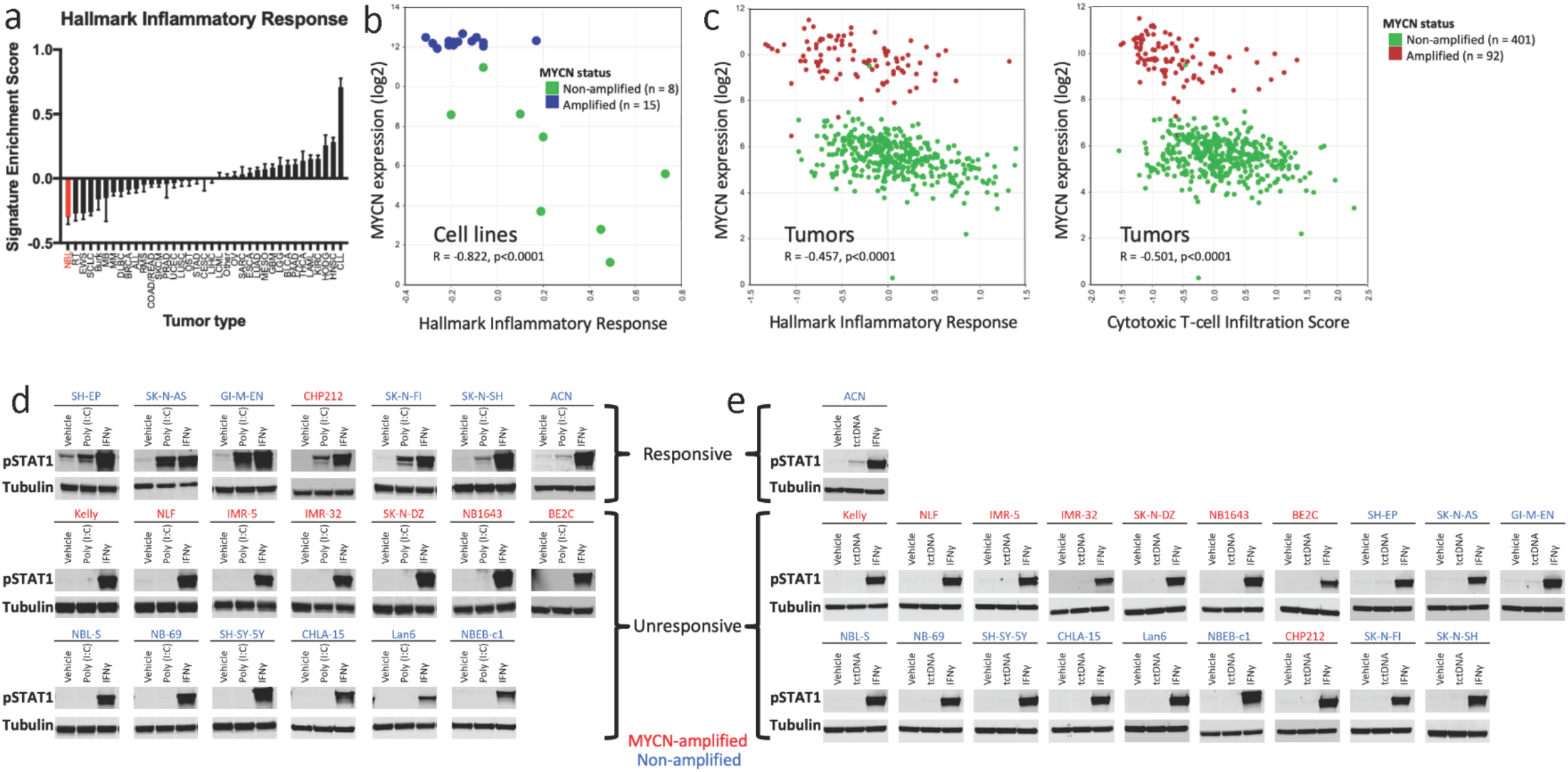
Neuroblastoma cells lack basal inflammatory signaling and respond heterogeneously to inflammatory stimuli. a) Relative enrichment score of the Hallmark Inflammatory Response signature across cell lines in the CCLE, with neuroblastoma shown in red. See Figure S1 for abbreviations. Data downloaded from the Broad CCLE portal (https://portals.broadinstitute.org/ccle). b) Comparison of *MYCN* expression and relative enrichment score of the Hallmark Inflammatory Response signature in 23 neuroblastoma cell lines. Data from <u>GSE28019</u>, obtained from and analyzed in R2 (http://r2.amc.nl). *MYCN-*amplified lines in blue, non-amplified in green. c) Comparison of *MYCN* expression and relative enrichment score of either the Hallmark Inflammatory Response signature (left) or a cytotoxic T cell infiltration score (right, from^77^) in 498 neuroblastoma tumors. Data from^78^, obtained from and analyzed in R2 (http://r2.amc.nl). *MYCN-*amplified tumors in red, non-amplified in green. d) Western blot showing differential changes in pSTAT1 levels when 20 neuroblastoma cell lines were treated with either 30μg/mL of poly (I:C) or 20ng/mL IFNγ for 24 hours. e) Western blot showing differential change in pSTAT1 levels when 20 neuroblastoma cell lines were treated with either transfected calf thymus DNA (tctDNA), IFNγ, or transfection reagent control (vehicle) for 6 hours. Western blots are representative of results from at least three separate experiments.

### Neuroblastoma cell lines respond heterogeneously to inflammatory stimuli

With the rationale that increasing cellular inflammatory signaling would increase HLA-I expression and lead to cytokine secretion that could drive T-cell recruitment, we next analyzed the response of a panel of 20 diverse neuroblastoma cell lines to different inflammatory stimuli (Figure 1d,e), including exogenous interferon gamma (IFNγ), poly (I:C) (polyinosinic:polycytidylic acid, a dsRNA-mimetic and TLR3 agonist), and transfected calf thymus DNA (tctDNA, agonist of cGAS). All of the cell lines tested generated a robust pSTAT1 response to exogenous IFNγ, consistent with previous reports of the effect of IFNγ on the expression of HLA-I in neuroblastoma cells^22, 41, 42^. Only one cell line responded strongly to the cGAS agonist, consistent with essentially undetectable expression of cGAS across neuroblastoma cell lines in the CCLE, in a different neuroblastoma RNA-seq data set^43^, and in RNA-seq data from neuroblastoma xenografts^44^ (Figure S2a-c). Similarly, western blot for cGAS in our cell line panel showed only the single responsive cell line had detectable expression (Figure S2d). Response to the TLR3 agonist, poly (I:C), however, was heterogenous, with the majority lacking response, while a subset showed robust responses. Intrigued by this heterogeneity, we decided to pursue the response to poly (I:C) further.

### TLR3-responsive cell lines demonstrate widespread inflammatory changes and increase susceptibility to T-cell killing

We next demonstrated in a subset of cell lines that the cells that responded to poly (I:C) with phosphorylation of STAT1 also responded by increasing transcription of IFN-response genes by qPCR, increasing nuclear localization of NF-ΚB, increasing phosphorylation of IRF3, and increasing activation of an NF-ΚB reporter construct (Figure S3a-c). Unresponsive cell lines showed none of these changes. We then performed transcriptomic analysis using 3’ RNA sequencing (Quantseq) on duplicate samples of 4 TLR3-responsive and 4 unresponsive cell lines treated with poly (I:C) or a vehicle control. Responsive cell lines showed a marked upregulation of inflammatory signaling pathways while unresponsive cell lines remained unchanged (Figure 2a and S4).

**Figure 2:**
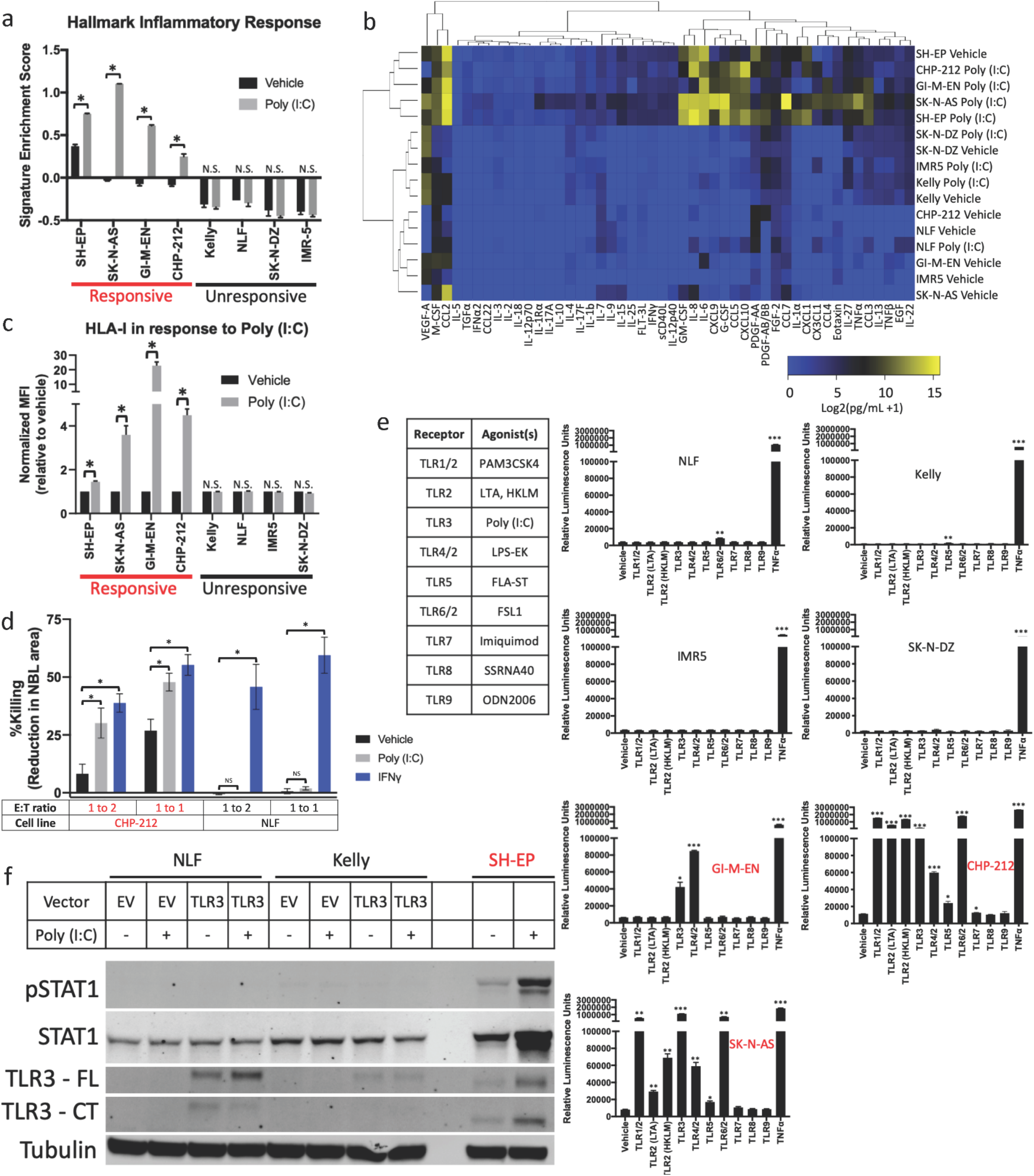
TLR3-responsive cell lines respond by multiple metrics and to additional TLR agonists. a) Relative enrichment of the Hallmark Inflammatory Response signature in the indicated cell lines treated with vehicle or 30μg/mL of poly (I:C) for 24 hours as measured by Quantseq. b) Heat map depicting the level of the indicated cytokines in the supernatant of the indicated neuroblastoma cell lines after treatment with vehicle or 30μg/mL of poly (I:C) for 24 hours. c) Change in surface expression of HLA-I, measured by flow cytometry, in the indicated neuroblastoma cell lines after treatment with vehicle or with 30μg/mL of poly (I:C) for 24 hours. d) Change in killing of neuroblastoma cells exogenously expressing MART-1 after culture with MART-1 transgenic tTCR-transfected T-cells. Neuroblastoma cells were treated with the indicated agonists for 24 hours then washed and cultured with the T-cells. E:T ratio is the effector (T-cell) to target (neuroblastoma) ratio. Killing was calculated by microscopy-based detection of change in cell area. e) Table showing the agonists used in the adjacent experiments. Relative luminescence in the indicated cell lines transfected with an NF-ΚB reporter after treatment with a vehicle control or the indicated TLR agonists for 6 hours. f) Western blot showing change in phosphorylation of STAT1, total STAT1, and TLR3 after treatment with 30μg/mL of poly (I:C) for 24 hours with or without expression of exogenous TLR3. Both the full length (FL) and an active C-terminal fragment (CT) of TLR3 are shown. TLR3-responsive cell lines are labelled in red, unresponsive in black. Two-tailed paired T-test between biological replicates, *p < 0.05, **p<0.01, ***p<0.001.

We then asked how poly (I:C) changed cytokine secretion, antigen presentation, and T-cell killing in neuroblastoma cell lines. We analyzed the concentration of 38 cytokines and chemokines in the supernatant of 4 responsive and 4 unresponsive cell lines cultured with and without poly (I:C). The profiles of the four responsive cell lines were similar once treated, while the unresponsive treated cell lines had profiles similar to the untreated lines (Figure 2b). A group of inflammatory cytokines were particularly increased in responsive cell lines when treated, including IL-6, IL-8, CCL5, CXCL9, CXCL10, G-CSF, and GM-CSF (Figure S5). Responsive cell lines also increased HLA-I surface expression as measured by flow cytometry, while unresponsive cell lines showed no change (Figure 2c), though all the cell lines responded to IFNγ (S3d). Lastly, we expressed MART-1 in one responsive (CHP-212) and one unresponsive (NLF) HLA-A2 neuroblastoma cell line (Figure S3e) and co-cultured them with T-cells transfected with a TCR targeting MART-1^45^. We found that the responsive cells were killed more easily by T-cells even prior to stimulation, and killing was enhanced by treatment with either poly (I:C) or IFNγ. In contrast, the unresponsive cells were resistant at baseline and killing was enhanced only by IFNγ, not by poly (I:C) (Figure 2d, S3f). Hence, TLR3 signaling can only be activated in a subset of neuroblastoma cells lines; in those lines it drives an inflammatory transcriptome, increases antigen presentation, and increases T-cell killing of neuroblastoma cells.

### Cell lines with active TLR3 sensing respond to additional TLR agonists

We next asked whether the defect in TLR3 signaling in unresponsive cell lines was limited to the expression of TLR3 itself, or whether this was indicative of a broader defect in inflammatory sensing in these cells that results in a widely insensitive state. We used an NF-ΚB reporter assay to test the response to 9 additional TLR agonists in three responsive cell lines and four unresponsive cell lines. While the responsive cell lines were also able to respond strongly to some additional stimuli (though heterogenous), the unresponsive lines did not show any strong responses (Figure 2e). Further supporting a more globally unresponsive state, we found that while unresponsive cell lines lacked TLR3 protein expression (Figure S2c), exogenous re-expression of TLR3 in unresponsive cell lines did not result in a response to poly (I:C), despite restoring TLR3 protein to a level similar to that found in a responsive cell line (Figure 2f). Together, these data pointed toward a state broadly unresponsive to TLR signaling rather than a single defect in TLR3 expression.

### MYCN plays a minor role in TLR3 response

Our first hypothesis for the driver of this insensitive state was *MYCN* amplification status. Amplified tumors have distinct transcriptomes^46^ and, as discussed, have lower markers of immune infiltration and inflammatory signaling^23, 24, 25, 26^. While 6 non-amplified lines responded to poly (I:C) and 6 others did not, only 1 of 8 amplified lines (CHP-212) showed any response. However, as shown in Figure S6a, this amplified line does have the high level of *MYCN* transcript and the robust MYCN signature^47^ that is typical of *MYCN-*amplified cells lines. We tested whether knocking down *MYCN* in unresponsive amplified lines could restore response to poly (I:C). We used either an inducible shRNA in BE2(c) cells^48^ or siRNAs in Kelly and NLF cells and found no change in response (Figure 3a,b and Figure S6b). While we achieved >80% knockdown in all three settings, given the supraphysiologic levels of MYCN signaling in such amplified lines, it is difficult to ascertain whether this knockdown resulted in a sufficiently “low” MYCN state. Therefore, to further test this hypothesis, we used two responsive non-amplified lines stably expressing a MYCN-ER construct^49, 50^, which allowed acute activation of MYCN signaling upon application of an estrogen receptor agonist. As shown in Figures 3c and 3d, acute MYCN activation only modestly decreased responsiveness to poly (I:C) and did not result in an insensitive state. Hence, we surmise that MYCN expression alone is not sufficient to explain the difference between responsive and unresponsive cell lines.

**Figure 3:**
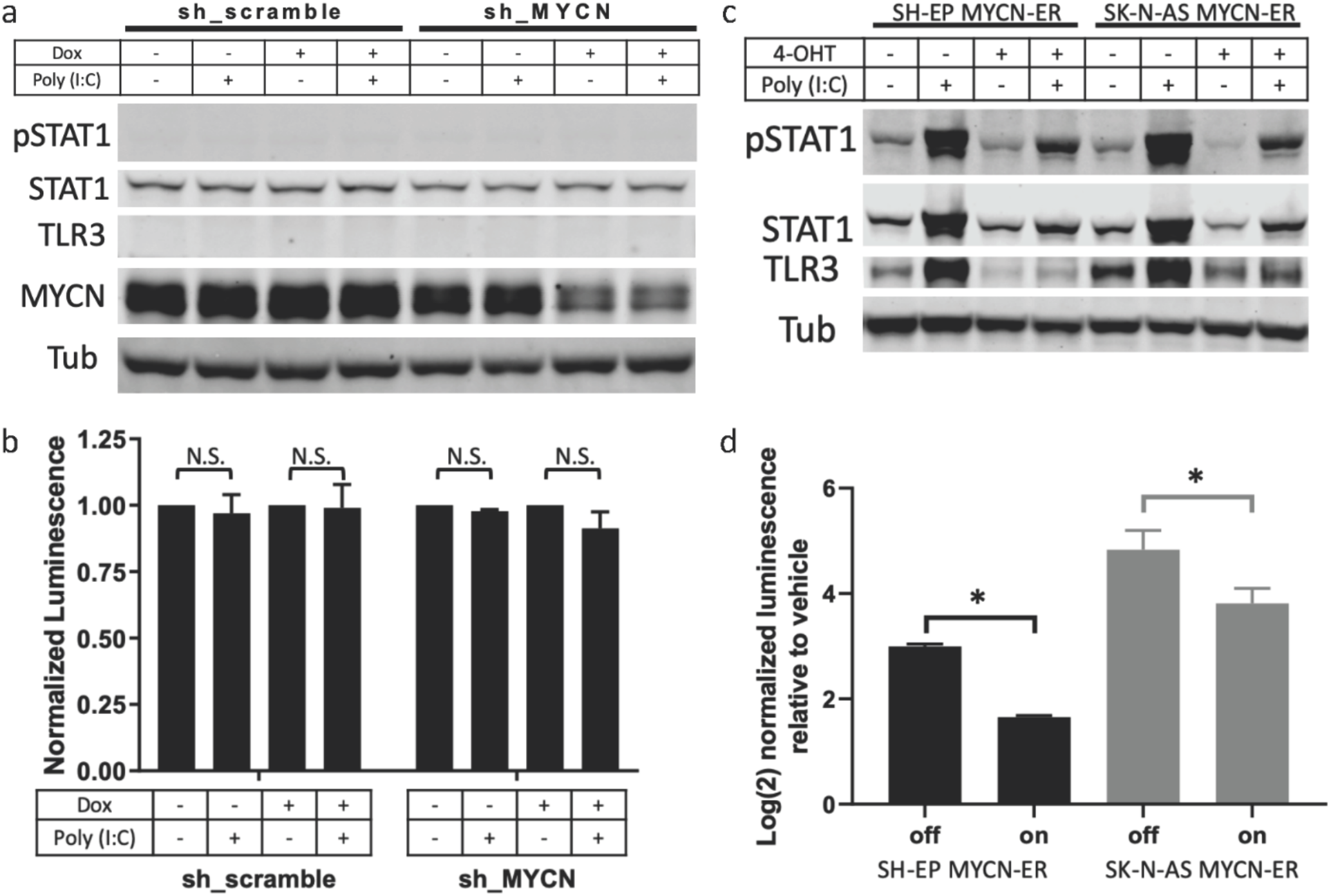
MYCN is insufficient to fully change TLR3-sensing state. a) Western blot showing changes in pSTAT1, STAT1, TLR3, and MYCN when BE2(c) cells expressing an inducible shRNA, either a scramble control or targeting MYCN, were treated with either vehicle or doxycycline (dox) for 72 hours, at which point they were then treated with either vehicle or 30μg/mL of poly (I:C) for 24 hours (with continued dox or vehicle). b) Change in luminescence detected after the cells used in panel (a) were transfected with an NF-ΚB reporter and then treated as described in (a). (c) Western blot showing changes in pSTAT1, STAT1, and TLR3 when SH-EP or SK-N-AS cells expressing a MYCN-ER construct were treated with vehicle or 4-hydroxytamoxifen (4-OHT) for 24 hours, at which point they were treated with either vehicle or 30μg/mL of poly (I:C) for 24 hours (with continued 4-OHT or vehicle). d) Change in luminescence detected after the cells used in panel (c) were transfected with an NF-ΚB reporter and then treated as described in (c). Each bar shows the increase comparing vehicle to poly (I:C) treatment in the specified cell line and MYCN-ER on/off condition. Western blots are representative of results from at least three separate experiments. Two-tailed paired T-test between biological replicates, *p < 0.001.

### MES vs ADRN status fully controls TLR3 response

Given the recently described MES and ADRN states of neuroblastoma cells^33, 34, 35^, we next hypothesized that these epigenetic states were associated with poly (I:C) response. Based on published categorizations of our cell lines^33, 34^ supplemented with additional RNA-seq data^43^ for unclassified lines (Figure S7a), we found that these states were highly predictive of poly (I:C) responsiveness (Figure 4a), strongly correlated with expression of TLR3 itself in cell lines and tumors (S7b), and predictive of H3K27 acetylation near the TLR3 promoter (Figure S7c). All of the unresponsive cell lines were ADRN, while all but one responsive cell line was MES. The SK-N-SH line was responsive. Interestingly, SK-N-SH was subcloned into two morphologically distinct lines, SH-EP and SH-SY5Y^51^, which have subsequently been classified as MES and ADRN, respectively^33^. Concordantly, SK-N-SH is responsive to poly (I:C), though not as strongly as SH-EP (the MES subclone), and SH-SY5Y (the ADRN subclone) is unresponsive. Lastly, the only *MYCN*-amplified line that responds to poly (I:C) is the only *MYCN*-amplified line in the MES state.

**Figure 4:**
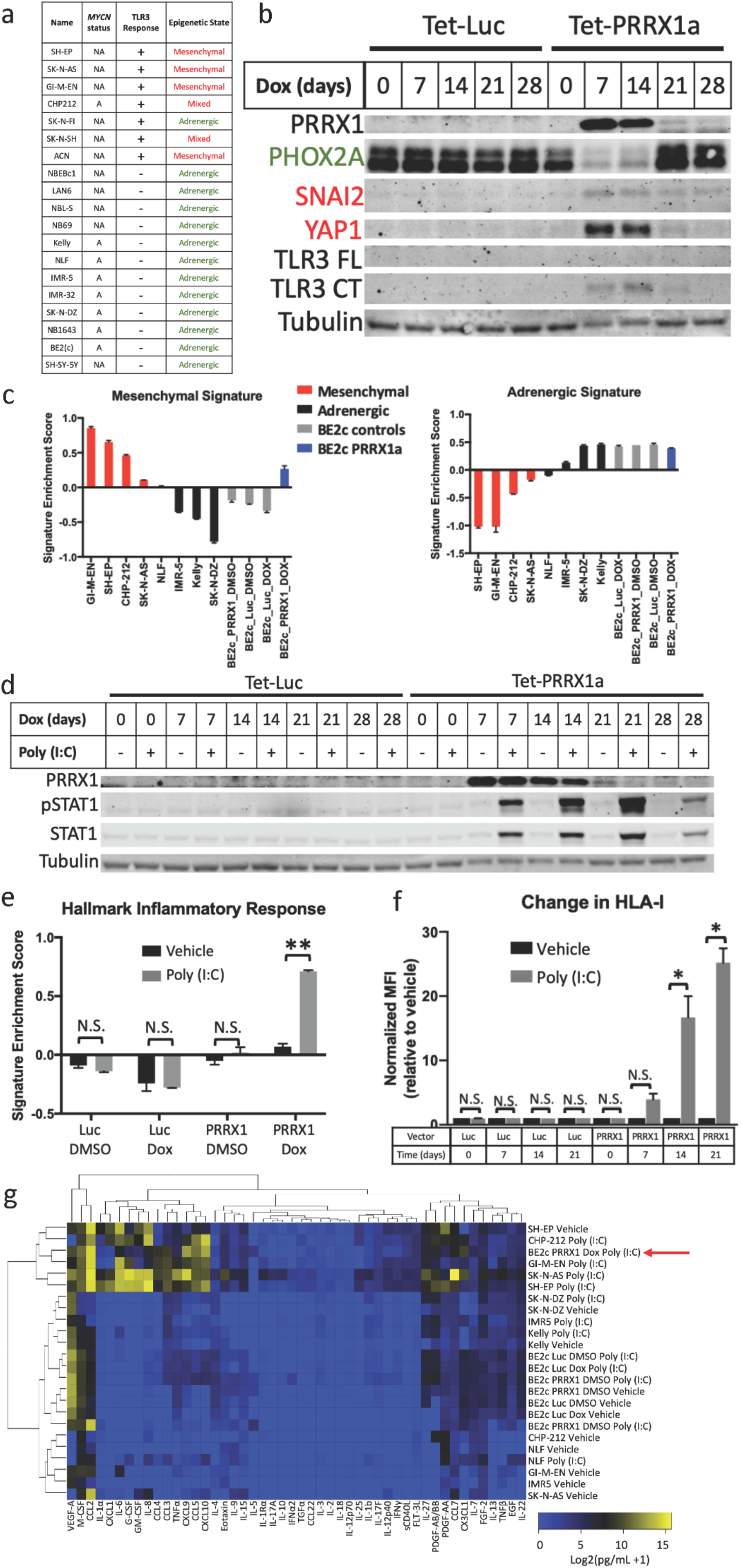
Epigenetic state determines TLR3-responsiveness. a) Table showing the cell lines used, their *MYCN* amplification status (A = amplified, NA = non-amplified), their TLR3 response, and their epigenetic state. Epigenetic state was determined based on prior reports (SH-EP, SK-N-AS, GI-M-EN, ACN, SH-SY-5Y, IMR32, BE2c, SK-N-FI from^33^, CHP-212, SK-N-SH, NB-69, NBEBc1, and SK-N-DZ from^34^) or from an expression signature-based comparison with characterized lines as shown in figure S7a. b) Western blot showing changes in the adrenergic marker PHOX2A, the mesenchymal markers SNAI2 and YAP1, and TLR3 after inducible expression of Luciferase (Luc) or PRRX1 for the indicated number of days. Both the full length (FL) and an active C-terminal fragment (CT) of TLR3 are shown. c) Relative enrichment the mesenchymal (left) and adrenergic (right) expression signatures in the indicated cell lines as measured by Quantseq. Inducible BE2(c) cells expressing either Luc or PRRX1 were. d) Western blot showing changes in pSTAT1 and total STAT1 when BE2(c) cells inducibly expressing either Luc or PRRX1 were treated with vehicle or doxycycline (Dox) for the indicated number of days, then treated with vehicle or 30μg/mL of poly (I:C) for 24 hours. Western blots are representative of results from at least three separate experiments. e) Relative enrichment of the Hallmark Inflammatory Response signature in BE2(c) cells expressing inducible Luc or PRRX1 treated with vehicle or doxycycline for 14 days, then treated with vehicle or 30μg/mL of poly (I:C) for 24 hours as measured by Quantseq. f) Change in surface expression of HLA-I, measured by flow cytometry, in BE2(c) cells expressing inducible Luc or PRRX1 treated with doxycycline for the indicated number of days, then treated with vehicle or with 30μg/mL of poly (I:C) for 24 hours. g) Heat map depicting the level of the indicated cytokines in the supernatant of the indicated neuroblastoma cell lines after treatment with vehicle or 30μg/mL of poly (I:C) for 24 hours. Inducible BE2(c) cells were treated with either vehicle (DMSO) or Doxycycline (Dox) for 14 days prior to poly (I:C) treatment. The red arrow highlights the BE2(c) cells with PRRX1 turned on and treated with poly (I:C). Western blots are representative of results from at least three separate experiments. Two-tailed paired T-test between biological replicates, *p<0.05, **p<0.02.

To determine whether the MES epigenetic state was simply correlated with TLR3-responsiveness or whether it was causally related, we sought to determine if altering the epigenetic state would also result in a change in TLR3-responsiveness. As has been described^33^, we inducibly expressed the mesenchymal transcription factor PRRX1a (or luciferase as a control) in the ADRN BE2(c) cell line. As shown in Figure 4b, this resulted in a loss of the adrenergic marker PHOX2A and new expression of the mesenchymal markers SNAI2 and YAP1. Quantseq data after a 14-day PRRX1a induction showed a strong increase in the MES signature score^33^, to a level comparable to native MES lines (Figure 4d), though only a modest decrease in the ADRN score. This switch also resulted in TLR3 protein expression (Figure 4b) and a stark conversion from unresponsive to TLR3 agonism to fully responsive (Figure 4d). Responsiveness measured by STAT1 phosphorylation was accompanied by upregulation of inflammatory expression signatures (Figures 4e and S8), secretion of inflammatory cytokines (Figures 4g, S9), and increase in HLA-I surface expression (Figure 4f), all of which were absent in the cell line without induction of transgene expression or expressing a luciferase control.

### MES vs ADRN status controls additional sensing pathways and basal inflammatory state

While these findings strongly suggested that cells in the MES state are responsive to stimulation of TLR3, we next asked whether the MES state may be associated with the broader active inflammatory sensing state that we found to be present in TLR3-responsive cell lines. The first evidence to suggest this is the case is that PRRX1a expression led to full restoration of the pSTAT1 response to poly (I:C), unlike exogenous TLR3 expression in an insensitive line, which did not lead to a response (Figure 2f and 4d). We next looked at the correlation between the MES gene signature^33^ and the transcripts of receptors, adaptors/signaling proteins, and effector transcription factors involved in TLR and other pattern recognition receptor signaling, in addition to how these transcripts changed with PRRX1 expression. In bulk tumors, nearly all of the examined transcripts were highly correlated with the MES signature, though this could be confounded by expression in stromal and immune cells. However, in addition to TLR3, several other transcripts were also highly correlated in two different cell line transcriptomic datasets (Table S1 and Figure S10a) and also increased with PRRX1a expression in our cells (Figure S10b). This included STING, which is activated by cGAS signaling, and the NF-ΚB subunit RELB, as well as TLR4 and TLR6. TLR4 is notably the receptor for the ligand LPS that was active in all three mesenchymal cell lines that were tested for response (Figure 2e). Concordantly, PRRX1 expression in BE2(c) cells resulted in a new sensitivity to LPS (Figure 5a). We next examined the relationship between MES and ADRN signatures and the basal inflammatory state in neuroblastoma by looking at correlations with inflammatory signaling genesets. In tumors, which are thought to be a heterogenous mix of the two states, we found that the MES signature score was highly correlated with both an inflammatory signaling geneset and with a transcriptional signature of cytotoxic T-cell infiltration (Figure 5b). This was also true for inflammatory signaling in cell lines (Figure 5c) and the reverse was true for the ADRN signature. PRRX1a expression in BE2(c) cells resulted in an increase in the inflammatory geneset to a level indistinguishable from the MES lines (Figure 5d). These strong correlations were not driven by overlapping genes in the genesets; for example, of the 200 total genes in the Hallmark Inflammatory Signaling geneset, 5 are in the 485 gene MES gene signature and 3 in the 369 gene ADRN gene signature. There is no overlap between the genes in the MES and ADRN signature and the cytotoxic T-cell infiltration signature. Hence, the MES state not only has activated TLR signaling, but also shows activation of additional immune sensing pathways and has an elevated basal inflammatory state.

**Figure 5:**
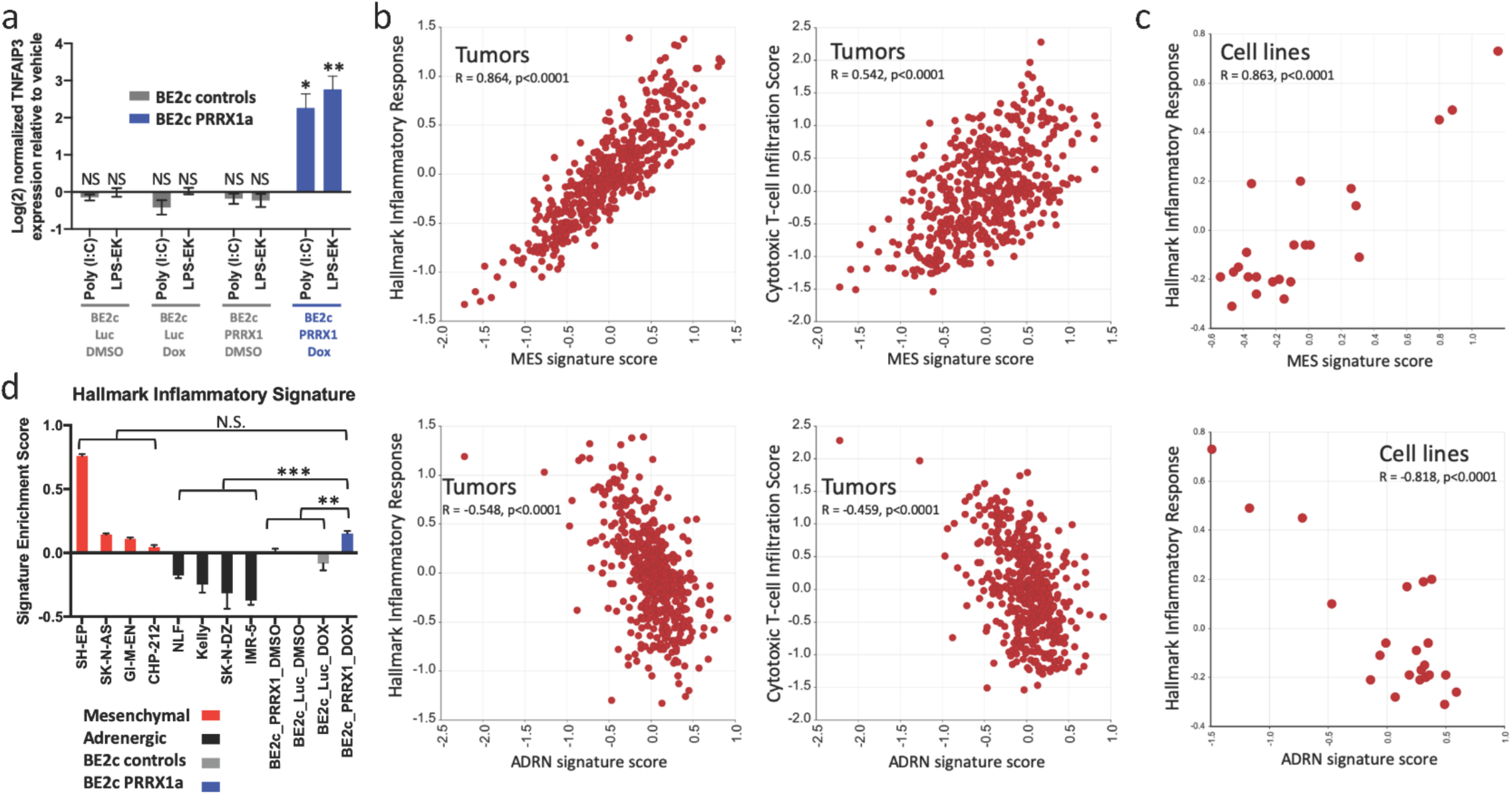
Epigenetic state controls additional TLRs and correlates with basal inflammatory state. a) Change in expression of *TNFAIP3* as measured by qPCR after treatment with the indicated agonists for 6 hours in BE2(c) cells after treatment with vehicle or doxycycline for 14 days. b) Comparison of the MES or ADRN signature scores^33^ to the enrichment of the Hallmark Inflammatory Response signature and to a cytotoxic T-cell infiltration signature^77^ in 498 neuroblastoma tumors^78^. Data obtained from and analyzed in R2 (http://r2.amc.nl). c) Comparison of the MES or ADRN signature scores to the enrichment of the Hallmark Inflammatory Response signature in 23 neuroblastoma cell lines (data from GSE28019). Data obtained from and analyzed in R2 (http://r2.amc.nl). d) Relative enrichment of the Hallmark Inflammatory Response signature in the indicated neuroblastoma cell lines as measured by Quantseq. BE2(c) cells expressing inducible luciferase control (Luc) or PRRX1 were treated with vehicle or doxycycline for 14 days. Two-tailed t-test between indicated groups, *p<0.05, **p<0.01, ***p<0.001.

### scRNAseq shows MES signature in human tumor cells correlates with inflammatory signature in vivo

Lastly, we asked whether MES/ADRN states are associated with inflammatory signaling in neuroblastoma cells *in vivo*. To address this, we analyzed single cell RNAseq data from 10 untreated adrenal high-risk neuroblastomas that were recently published^52^. Classification by cell identity marker genes demonstrated that the majority of cells showed a neuroendocrine phenotype with smaller numbers of immune cells, Schwann cells, and stromal cells (Figure 6a). We then used the same approach taken by the authors of this study to infer chromosomal copy number variations (CNVs) from the transcriptomic data individually for each sample, which allowed for separation of presumed tumor and non-tumor cells (Figure S11a). Similar to the original study, we did not identify a transcriptionally distinct population of pure tumor cells with mesenchymal markers. However, there is substantial transcriptional variability within the tumor compartment, including within individual tumors (Figure S11a). We therefore asked whether cells with stronger ADRN or MES signatures could be identified. Indeed, the two signatures were negatively correlated overall and in every tumor (Figures 6b, S11b). We then compared the MES signature to the Hallmark Inflammatory Signature and found that these signatures were strongly positively correlated overall and in every individual tumor (Figures 6c, S11c). If the cells with the highest MES score and lowest ADRN score are compared to the cells with the lowest MES and highest ADRN score, the difference in inflammatory signature is even more striking (Figure 6d). This demonstrates that in human neuroblastoma tumors, cells with stronger MES signatures have higher levels of basal inflammatory transcripts compared to those with stronger ADRN signatures. Similar to cell lines with higher MES character, we postulate that these cells are thus likely to be uniquely vulnerable to inflammatory stimuli and therefore may be more susceptible to immune-based therapies. Given the reports of chemotherapy resistance in this subset of cells, immune targeting of these cells has the potential to improve outcomes and decrease relapse in neuroblastoma.

**Figure 6:**
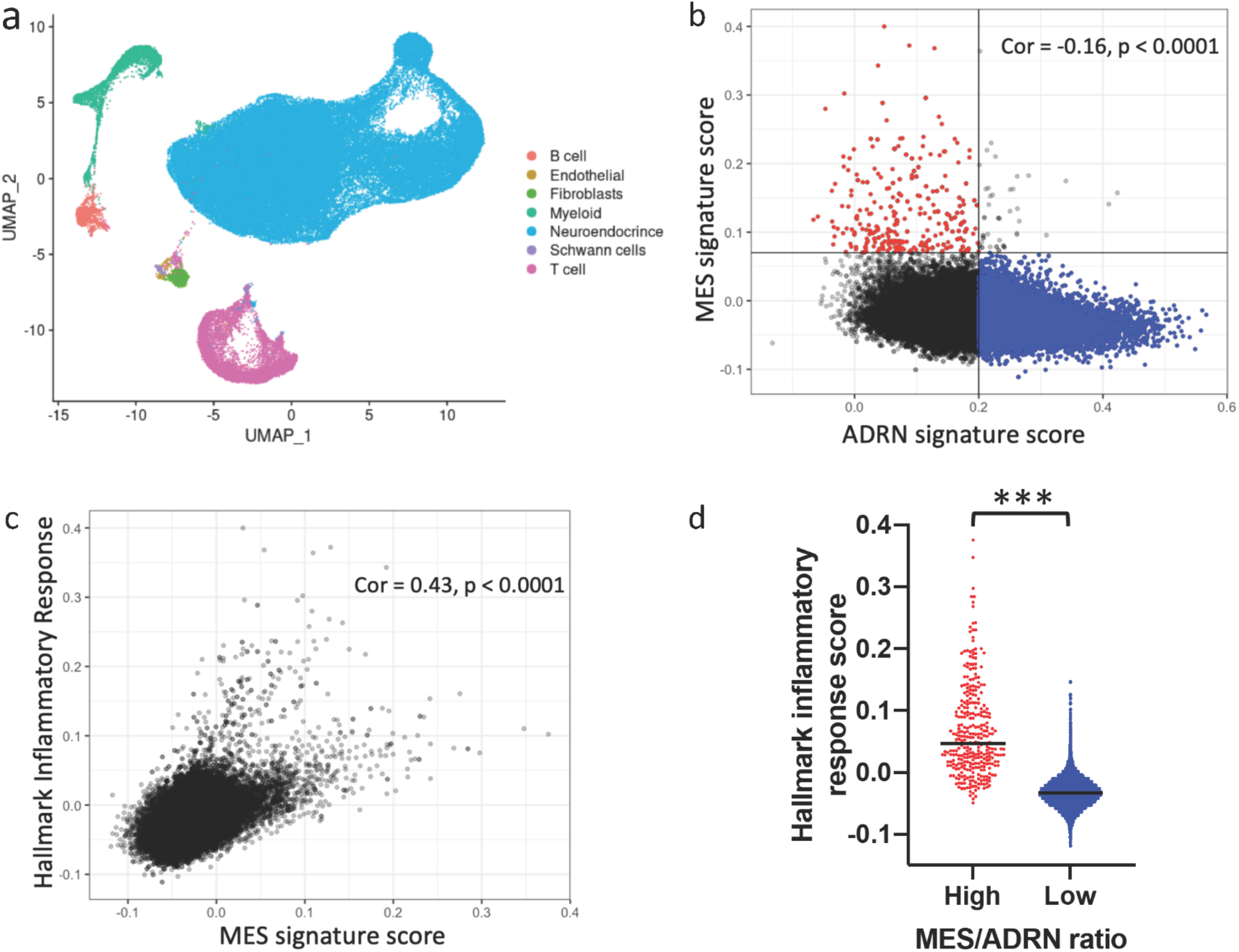
scRNAseq in human neuroblastoma shows inflammatory signature correlates with MES signature. a) UMAP plot showing cell type distributions in the 10 pooled samples. b) Correlation between the MES and ADRN gene signatures^33^ in the cells determined to be tumor based on CNV. Cells are shown from all 10 samples. Quadrant in red is defined as high MES/ADRN ratio, quadrant in blue is defined as low MES/ADRN ratio. c) Correlation between the Hallmark Inflammatory Response signature^39^ and the MES signature in the cells determined to be tumor based on CNV. Cells are shown from all 10 samples. d) Graph depicting the Hallmark Inflammatory Signature score for each cell with high MES/ADRN ratio (red) vs low MES/ADRN ratio (blue) as defined in panel (b). Black bar indicates the mean. Two-tailed t-test between indicated groups, ***p<0.001.

## Discussion

Immunotherapy has had a transformative effect on cancer therapy. This includes neuroblastoma, as antibodies against GD2 improves survival when combined with immune-stimulating cytokines, although cure remains elusive in large numbers of patients. Other immunotherapies, such as checkpoint inhibitors and CAR T cells, have had limited efficacy. Here we show that neuroblastoma is a particularly difficult challenge, even among cold tumors. Using cell line transcriptomic data, we found that neuroblastoma cells are the most deficient in inflammatory signaling among the 37 cancer histologies examined. While such analysis will be more definitively performed once widespread single cell RNA-seq data is available across tumor types, the current data provides further evidence that neuroblastoma cells lack substantial inflammatory signaling at baseline. However, there is reason for optimism that altering that cold state may allow improved entry, persistence, and activity of activated immune cells^6, 7^. As a precursor to enhancing immunotherapy in neuroblastoma, we investigated the pathways that could be stimulated in order to activate inflammatory signaling in neuroblastoma cells.

Multiple prior studies have shown that neuroblastoma cell lines increase HLA-I upon stimulation with IFNγ^22, 41, 42^. Consistent with this finding, we found that all cell lines generated an increase in pSTAT1 upon treatment with IFNγ. A prior clinical study treated neuroblastoma patients with IFNγ^53, 54^, and a more recent analysis^42^ showed that this increased T cell infiltration and HLA-I expression in human tumors and had similar effect in mouse models. This study was done prior to the development of checkpoint inhibitors, but such an approach could be revisited. Only one of the 20 cell lines tested responded to cGAS stimulation or had any detectable expression of cGAS. Sensing by cGAS has been implicated in driving inflammatory responses to chromothripsis^11^, an important process in poor-prognosis neuroblastoma^55^. Downregulation of cGAS may explain the ability of neuroblastoma to avoid the immune-stimulating effects of chromothripsis or of extra-chromosomal circular DNA, which is found widely in neuroblastoma including as a frequent mechanism of *MYCN* amplification^56^. Given the essential role of cGAS-STING signaling in mediating a productive interaction between radiation therapy and ICB^15^, our findings suggest that radiation therapy could be optimized by strategies to activate cGAS-STING in neuroblastoma to improve the systemic immune response.

We focused our work on exploring and understanding the heterogenous responses of neuroblastoma cells to TLR3 agonism. Response to TLR3 agonists have previously been explored in neuroblastoma, with most studies focused on activation of cell death^57, 58, 59, 60^. Rather than induction of cell death, we concentrated on activation of inflammatory signaling, as this was our goal and since we saw little to no death after short treatments (<48 hours, data not shown). A prior study in neuroblastoma showed that in some cell lines, TLR3 stimulation can increase PD-L1 and MHC-I surface expression, increase secretion of inflammatory cytokines, and improve activation of T-cells^61^. Consistent with this, we found that, only the subset of responsive cell lines, treatment with a TLR3 agonist resulted in multiple changes associated with an improved immune response. This included activation of an inflammatory transcriptome, secretion of immune-recruiting cytokines, and increase in antigen presentation. Concordantly, we were able to show that TLR3 activation increases the killing of neuroblastoma cell by T cells only in responsive lines.

Prior studies in neuroblastoma had noted some differential responses to poly (I:C) and differences in TLR3 expression^58, 59, 60, 61^, but no mechanistic basis for this difference had been identified. In retrospect, the responsive cell lines in these studies are from the MES subtype, consistent with our results. The use of a much larger cell line panel, together with the more recent descriptions of super-enhancer-associated epigenetic states, allowed us to connect these states to TLR3 signaling. By showing that inducing a shift toward the MES state in an unresponsive cell line is sufficient to convert it to a fully responsive state, we show epigenetic state is more important than *MYCN* amplification status and convincingly connect these epigenetic states to inflammatory sensing. This connection had not been previously made, to our knowledge. Questions remain about the specific nature of this link, whether it is mediated by a single or multiple transcription factors, and how inextricable the epigenetic state is from immune sensing.

Using a published scRNAseq dataset^52^, we were able to show that tumor cells with stronger MES signatures have higher basal levels of inflammatory signaling, providing evidence supporting the hypothesis that MES cells *in vivo* are likely to be more responsive to inflammatory stimuli and more sensitive to immunotherapies. Similar to the findings of the group that originally published this dataset, we did not identify a clear and separate population of tumor cells with mesenchymal markers, but there is substantial intra- and inter-tumoral transcriptional heterogeneity. As this dataset is composed exclusively of untreated adrenal primary tumors, it may be that MES populations become more distinct after therapy or at relapse. Alternatively, it may be that cells are in a continuous distribution between the MES/ADRN states and that their cell identity is not as binary *in vivo* as cell lines tend to be *in vitro*.

The optimal approach to translating knowledge about the state of immune sensing pathway into therapeutic approaches remains an open question. As mentioned, IFNγ has been used in neuroblastoma patients in the past, though such a treatment has potential systemic drawbacks. STING agonists have generated significant interest across oncology^62^, including neuroblastoma^63^. A more stable analog of poly (I:C) was used systemically in neuroblastoma patients^64^, without significant efficacy and with marked systemic side effect, although a more recent study using a different dosing scheme showed little toxicity and some efficacy in children with brain tumors ^65^ and is currently being used widely in clinical trials. Our data suggests that an understanding of MES and ADRN states may contribute to the choice of the optimum immunostimulatory pathway to use in neuroblastoma and the most advantageous timing. For example, therapies that aim to target STING or TLRs may be more effective at later stages of therapy when responsive MES cells compose a higher percentage of remaining tumor cells. Indeed, the higher basal inflammatory state of residual MES cells may in part explain the effectiveness of anti-GD2 antibody immunotherapy when used in a minimal residual disease state^30^. Comparatively, there is a relatively poor response when used alone in the setting of bulk tumor^66^, which is likely to contain a significant portion of immunologically quiescent adrenergic cells. However, responses are much better when anti-GD2 therapy is combined with chemotherapy^31, 32^, which may allow for targeting of both states simultaneously.

Immunocompetent animal models will be needed to understand the impact of the differences in inflammatory sensing between MES and ADRN states *in vivo* and a lack of such experiments is a limitation of the current study. This provides a challenge, however. The most widely used neuroblastoma genetically engineered mouse model is the TH-MYCN model, with MYCN expressed from the tyrosine hydroxylase promoter^67^. While there is some evidence that cells from these tumors can assume a mesenchymal-like state in the setting of treatment resistance^37^, tyrosine hydroxylase expression is much lower in the MES state^36^, which in the TH-MYCN model may turn off the expression of MYCN. Further work is first needed to explore the MES and ADRN states in the TH-MYCN and other lesser-used neuroblastoma models.

In conclusion, here we have shown here that neuroblastoma cells have uniquely suppressed inflammatory signaling and assessed approaches to modify that signaling. We found that super-enhancer-associated epigenetic states determine the activity of TLR pathway sensing, which, when activated, can lead to a host of alterations expected to improve immune detection of tumors. This work demonstrates a new role for intratumoral heterogeneity in modifying inflammatory signaling. Given the distinct transcriptional and regulatory networks of MES and ADRN cells and the resistance of the MES state to cytotoxic chemotherapy, it is likely that curing neuroblastoma will require simultaneous targeting of both states with separate, but perhaps synergistic, therapies. Our data suggest that immune-directed therapies may be able to exploit a specific vulnerability of the MES state. Further work is needed to therapeutically exploit this vulnerability.

## Methods

### Cell Culture

Cell lines were generous gifts from Drs. Marie Arsenian-Henriksson, Michael Hogarty, John Maris, Michael Milone, and Linda Valentijn or purchased from either the German Collection of Microorganisms and Cell Cultures or the Interlab Cell Line Collection (see Supplementary Table 2 for specific details). Cells were maintained at 37 °C and 5% CO_2_ in a humidified incubator and cultured with media components listed. Cell identities were verified by STR profiling and routinely tested negative for mycoplasma contamination.

### Reagents

Poly (I:C) was purchased from Invivogen (tlrl-pic) and used at 30 μg/mL. Recombinant IFNγ was purchased from Peprotech (300-02) and used at 20 ng/mL. PAM3CSK4 (1 μg/mL), LTA-SA (10 μg/mL), FLA-ST (100 ng/mL), FSL-1 (4 μg/mL), Imiquimod (5 μg/mL), ssRNA40 (5 μg/mL), LPS-EK (1 μg/mL) and ODN2006 (10 μg/mL) were purchased from InvivoGen (tlrl-kit1hw and tlrl-pslta). Calf thymus DNA was purchased from Sigma (D1501), used 1ug/mL, and transfected with jetPRIME transfection reagent (VWR 89129-920). Doxycycline (Sigma D9891) was used at 50 ng/mL for inducible shRNA experiments and 1μ/mL for PRRX1a overexpression. Qiagen FlexiTube MYCN siRNA (SI03113670, SI03087518) and negative control (1027280), 30 pmol/well, were transfected using Lipofectamine^™^ RNAiMax Transfection reagent (13778150) according to the manufacturer’s protocol. For MYCN-activation experiments, 4-hydroxytamoxifen (Sigma H7904) was used at 500nM. The concentration for drug selection using zeocin was determined for each cell line independently. Puromycin was used at 2.5 μg/mL for BE2(c) cells and 1μg/mL for CHP-212 and NLF cells. G418 was used at 500 μg/mL for Kelly cells and 250 μg/mL for NLF cells. The Luciferase Assay System (E1501) was purchased from Promega and used according to the manufacturer’s instructions.

### Plasmids

NF-κB pNifty2-Luc was purchased from InvivoGen and transfected into cell lines using Lipofectamine^™^ 3000 transfection reagent (L3000008) followed by zeocin selection. Myc-DDK-PRRX1a and Myc-DDK-TLR3 were purchased from Origene (RC213276, RC210497). TLR3 was transfected using Lipofectamine^™^ 3000 and selected with G418. PRRX1a was cloned into the pLVX-TetOne (Takara 631847). Final plasmid identity was validated by sequencing. Lentivirus was produced from the PRRX1a plasmid and the analogous luciferase control and cells were infected as previously described^68^. Infected cells were selected with puromycin. Lentiviral particles for MART-1 expressed from an EF1a promoter with an IRES-eGFP were purchased from Genecopoeia (G0616-Lv225). GFP control was expressed using a lentiviral vector with an EF1a promoter and IRES-eGFP.

### Cytokine profiling

Cytokine profiling was performed by the Human Immunology Core at the University of Pennsylvania using the Luminex FlexMap 3D system (EMD Millipore). A MILLIPLEX human cytokine 48-plex panel (Millipore Sigma HCYTA-60K-PX48) was run using supernatant from cells cultured in the presence or absence of poly I:C for 24 hours. Samples were run in biological triplicate. Cytokine concentrations were obtained by comparison to a standard curve.

### Western blots

Cells were lysed on ice using M-PER (Thermo Scientific 78501) with phosphatase inhibitor cocktails 2 and 3 (Sigma Aldrich P5726, P0044) and a protease inhibitor cocktail from Promega (G6521) then scraped and cleared by centrifugation at 4°C. Protein was quantified using the DC Protein Assay (Bio-Rad) then electrophoresed using Criterion TGX gels (Bio-Rad) and transferred to nitrocellulose membranes via iBlot2. Membranes were blocked in Odyssey blocking buffer. The following primary antibodies were used for immunoblotting, diluted in TBS-T with 5% bovine serum albumin: rabbit phospho-Tyr701 STAT1 (Cell Signaling 9167), rabbit STAT1 (Cell Signaling, 9172S), mouse PHOX2A (Santa Cruz 81978), mouse PRRX1 (Origene TA803116), rabbit YAP (Cell Signaling 14074), mouse TLR3 (Novus Biologicals NBP2-24875), rabbit SLUG (SNAI2, Cell Signaling 9585), rabbit STING (Cell Signaling 13647), mouse α-tubulin (Calbiochem CP06), rabbit α-tubulin (Cell Signaling 2144). Secondary antibodies included goat anti-rabbit Alexa Fluor 680 (Invitrogen A21109), goat anti-mouse Alexa Fluor 790 (Invitrogen A11357), and goat anti mouse DyLIght^™^ 800 (5257). Immunoblots were visualized by an Odyssey CLx (LI-COR).

### Flow cytometry

Cells were trypsinized and resuspended in PBS with 2% FBS at 1×10^6 cells/mL, then incubated with antibodies (1:100) at 37*C for 30 minutes. Cells were washed twice with 2% FBS in PBS, then resuspended in 50% accutase (Sigma A6964) and 50% of PBS with 2% FBS. Flow cytometry was executed using a Guava EasyCyte flow cytometer (Luminex). PE-HLA-A,B,C antibody (311406) and isotype control (400214) were purchased from BioLegend. Data were analyzed in FlowJo.

### T-cell killing assay

Primary human T-cells were collected, transfected, and expanded as previously described^45^ and stored as frozen aliquots on day 13 after bead activation. T-cells were thawed, rested for one day, and activated with ImmunoCult™ Human CD3/CD28 T Cell Activator (Stem Cell Technologies) following the manufacturer’s protocol. T-cell cultures were fed with fresh 10% RPMI supplemented with human IL2 (20u/ml Stem Cell Technologies) every 2-3 days to maintain 1×106 cells/mL/mm2. T-cells were rested for a minimum of 11 days before use in functional assays. Neuroblastoma cells were seeded in 96-well plates then treated with vehicle, 30 μg/mL poly (I:C), or 20 ng/mL IFNγ for 24 hours. Cells were then washed with media and T-cells were added at the indicated effector to target ratio. After 18 hours, 12 non-overlapping fields per well of 96-well plate were captured at 10X using a Nikon Ti-E Automated Inverted Microscope with Nis-Elements Ar. Nis Jobs template “Fixed Cells” was used to define image acquisition parameters. Tif images were processed and converted to Simple Segmentation images using in the Ilastik 1.3.3 (interactive machine learning for (bio)image analysis) tool^69^. The Simple Segmentation images were analyzed for GFP area in CellProfiler 4.07 (cell image analysis software)^70^ and exported into Microsoft Excel.

### Quantitative PCR and Quantseq

For both Quantseq and qPCR, RNA was extracted using the Qiagen RNeasy Plus mini kit (74134) after homogenization by centrifugation in QIAShredder (79656) microcentrifuge tubes. For qPCR, RNA was reverse transcribed with a TaqMan Reverse Transcription Reagent Kit (Applied Biosystems N8080234, using oligo(dT)). Relative transcript abundance was obtained using Power SYBR Green Master Mix (ThermoFisher) using the QuantStudio 6 Flex Real-Time PCR System using primers for OAS1 (GATGAGCTTGACATAGATTTGGG and GGTGGAGTTCGATGTGCTG) and TNFAIP3 (CAAGTGGAACAGCTCGGATT and GCCCAGGAATGCTACAGATAC) as well as RPLP0 (TGTCTGCTCCCACAATGAAAC and TCGTCTTTAAACCCTGCGTG) as a normalization control. For Quantseq, RNA quantity was determined using the Qubit 2.0 Fluorometer (ThermoFisher Scientific, Waltham, MA) and the quality was validated using the TapeStation RNA ScreenTape (Agilent, Santa Clara, CA). 200 ng of DNAse I treated, total RNA was used to prepare library for Illumina Sequencing using the Quant-Seq 3’mRNA-Seq Library Preparation Kit (Lexogen, Vienna, Austria). Library quantity was determined using qPCR (KAPA Biosystem, Wilmington, MA). Overall library size was determined using the Agilent TapeStation and the DNA High Sensitivity D5000 ScreenTape (Agilent, Santa Clara, CA). Equimolar amounts of each sample library were pooled, denatured and sequenced using High-Output, Single-read, 75bp cycle Illumina NextSeq 500 sequencing kits. Next Generation Sequencing was done on a NextSeq 500 (Illumina, San Diego, CA).

### Database and Quantseq analysis

Data was downloaded from the Broad CCLE portal (https://portals.broadinstitute.org/ccle), from <u>GSE89413</u>^43^, or PedcBioPortal (https://pedcbioportal.kidsfirstdrc.org)^44^ as FPKM values, then converted to log_2_(FPKM+1). For Quantseq, raw RNA-seq reads were trimmed using Trimmomatic v.0.39^71^ and data was aligned using the bowtie2^72^algorithm against hg38 human genome version. RSEM v1.2.12 software^73^ was used to estimate read counts using gene information from Ensemble transcriptome version GRCh37.p13. Mesenchymal and adrenergic genesets were taken from van Groningen and colleagues^33^. Other genesets were downloaded from the Molecular Signatures Database (www.gsea-msigdb.org). Each gene in each signature was z-transformed across the CCLE cell lines and the signature score calculated as the average of the z-transformed gene scores. Other datasets were analyzed with tools built into the R2: Genomics Analysis and Visualization Platform (http://r2.amc.nl).

### Single-cell RNAseq analysis

Count matrices for single cell analysis were downloaded from <u>GSE137804</u>. Non-high-risk patients were excluded from the analysis. Of note, sample T71 was excluded as the clinical description provided does not fit standard Children’s Oncology Group defined high-risk group status based on age, stage, and MYCN status. Gene expression matrices for each sample were analyzed using the Seurat package^74^. DoubletFinder^75^ was used to remove doublets. Cells with less than 500 features, 1000 counts, or greater than 10% mitochondrial counts were filtered out. The most variable genes were used to perform Principal Component Analysis (PCA) and the top 20 PCs were included to perform Louvain clustering. Cell types were annotated using the following gene markers: PHOX2B, DBH, TH, ISL1, GATA3 and HAND2 for neuroendocrine cells, S100B, SOX10 and PLP1 for Schwann cells, CD2, CD3D and CD247 for T-cells, CD79A, PAX5 and BANK1 for B-cells, CD14 and CD68 for myeloid cells, COL1A1, LAMA2 and COL3A1 for fibroblasts, and VWF, CALCRL and FLT1 for endothelial cells. In order to identify malignant cells, inferCNV^76^ was used. Specifically, inferCNV was applied for each sample using sympathoblasts, myeloid and fibroblasts from fetal adrenal of samples F6, F7, F106, F107 as a reference. To determine the ADRN, MES, and inflammatory score of each cell, the AddModuleScore function in the Seurat R package was applied using previously published signatures^33, 39^. We then calculated Pearson correlation between adrenergic and mesenchymal score vectors and between inflammatory and MES score vectors.

## Acknowledgments

The authors would like to thank Drs. Marie Arsenian-Henriksson, Michael Milone, and Linda Valentijn for providing cell lines used in the study and Dr. Zachary Stine and Rebekah Brooks for careful reading of the manuscript and helpful discussions. This work was supported by Alex’s Lemonade Stand Young Investigator Award (AJW), Hyundai Hope on Wheels Young Investigator Award (AJW), NIH K12 HD043245 (AJW), NIH K12 CA076931 (AJW), NIH R35 CA220500 (JMM), Giulio D’Angio Endowed Chair (JMM), NIH R01 CA057341 (CVD), NIH R01 CA051497 (CVD), and the Ludwig Institute for Cancer Research (CVD).

## Author Contributions

Conceptualization (AJW, MDH, JMM, CVD), Formal analysis (AJW, LDG, YU), Investigation (AJW, MMD, JLD, PAB, DV), Resources (ARG, DJP), Writing (original draft – AJW, review and editing – AJW, MDH, JMM, CVD), Supervision (AJW, DJP, KT, MDH, JMM, CVD), Funding acquisition (AJW, DJP, KT, MDH, JMM, CVD).

## Competing Interests Statement

CVD serves on the board of Rafael Pharmaceuticals, Inc. and as an advisor to the Barer Institute, Inc. The other authors declare no potential conflicts of interest.

## References

1. Tumeh PC, et al. PD-1 blockade induces responses by inhibiting adaptive immune resistance. Nature 515, 568–571 (2014).

2. Rizvi NA, et al. Cancer immunology. Mutational landscape determines sensitivity to PD-1 blockade in non-small cell lung cancer. Science 348, 124–128 (2015).

3. Yarchoan M, Hopkins A, Jaffee EM. Tumor Mutational Burden and Response Rate to PD- 1 Inhibition. N Engl J Med 377, 2500–2501 (2017).

4. Cristescu R, et al. Pan-tumor genomic biomarkers for PD-1 checkpoint blockade-based immunotherapy. Science 362, (2018).

5. Ayers M, et al. IFN-gamma-related mRNA profile predicts clinical response to PD-1 blockade. The Journal of clinical investigation 127, 2930–2940 (2017).

6. Zemek RM, et al. Sensitization to immune checkpoint blockade through activation of a STAT1/NK axis in the tumor microenvironment. Science translational medicine 11, (2019).

7. Ishizuka JJ, et al. Loss of ADAR1 in tumours overcomes resistance to immune checkpoint blockade. Nature 565, 43–48 (2019).

8. Junt T, Barchet W. Translating nucleic acid-sensing pathways into therapies. Nature reviews Immunology 15, 529–544 (2015).

9. Apetoh L, et al. Toll-like receptor 4-dependent contribution of the immune system to anticancer chemotherapy and radiotherapy. Nature medicine 13, 1050–1059 (2007).

10. Chiappinelli KB, et al. Inhibiting DNA Methylation Causes an Interferon Response in Cancer via dsRNA Including Endogenous Retroviruses. Cell 162, 974–986 (2015).

11. Mackenzie KJ, et al. cGAS surveillance of micronuclei links genome instability to innate immunity. Nature 548, 461–465 (2017).

12. Boelens MC, et al. Exosome transfer from stromal to breast cancer cells regulates therapy resistance pathways. Cell 159, 499–513 (2014).

13. Liu Y, et al. Tumor Exosomal RNAs Promote Lung Pre-metastatic Niche Formation by Activating Alveolar Epithelial TLR3 to Recruit Neutrophils. Cancer Cell 30, 243–256 (2016).

14. Sistigu A, et al. Cancer cell-autonomous contribution of type I interferon signaling to the efficacy of chemotherapy. Nature medicine 20, 1301–1309 (2014).

15. Harding SM, Benci JL, Irianto J, Discher DE, Minn AJ, Greenberg RA. Mitotic progression following DNA damage enables pattern recognition within micronuclei. Nature 548, 466–470 (2017).

16. Siegel RL, Miller KD, Jemal A. Cancer Statistics, 2017. CA Cancer J Clin 67, 7–30 (2017).

17. Maris JM. Recent advances in neuroblastoma. N Engl J Med 362, 2202–2211 (2010).

18. Brodeur GM, Seeger RC, Schwab M, Varmus HE, Bishop JM. Amplification of N-myc in untreated human neuroblastomas correlates with advanced disease stage. Science 224, 1121–1124 (1984).

19. Coughlin CM, et al. Immunosurveillance and survivin-specific T-cell immunity in children with high-risk neuroblastoma. J Clin Oncol 24, 5725–5734 (2006).

20. Pugh TJ, et al. The genetic landscape of high-risk neuroblastoma. Nature genetics 45, 279–284 (2013).

21. Wolfl M, et al. Expression of MHC class I, MHC class II, and cancer germline antigens in neuroblastoma. Cancer Immunol Immunother 54, 400–406 (2005).

22. Raffaghello L, et al. Multiple defects of the antigen-processing machinery components in human neuroblastoma: immunotherapeutic implications. Oncogene 24, 4634–4644 (2005).

23. Layer JP, et al. Amplification of N-Myc is associated with a T-cell-poor microenvironment in metastatic neuroblastoma restraining interferon pathway activity and chemokine expression. Oncoimmunology 6, e1320626 (2017).

24. Zhang P, et al. MYCN Amplification Is Associated with Repressed Cellular Immunity in Neuroblastoma: An In Silico Immunological Analysis of TARGET Database. Front Immunol 8, 1473 (2017).

25. Wei JS, et al. Clinically Relevant Cytotoxic Immune Cell Signatures and Clonal Expansion of T-Cell Receptors in High-Risk MYCN-Not-Amplified Human Neuroblastoma. Clinical cancer research : an official journal of the American Association for Cancer Research 24, 5673–5684 (2018).

26. Bernards R, Dessain SK, Weinberg RA. N-myc amplification causes down-modulation of MHC class I antigen expression in neuroblastoma. Cell 47, 667–674 (1986).

27. Davis KL, et al. Nivolumab in children and young adults with relapsed or refractory solid tumours or lymphoma (ADVL1412): a multicentre, open-label, single-arm, phase 1-2 trial. The Lancet Oncology, (2020).

28. Merchant MS, et al. Phase I Clinical Trial of Ipilimumab in Pediatric Patients with Advanced Solid Tumors. Clinical cancer research : an official journal of the American Association for Cancer Research 22, 1364–1370 (2016).

29. Richards RM, Sotillo E, Majzner RG. CAR T Cell Therapy for Neuroblastoma. Front Immunol 9, 2380 (2018).

30. Yu AL, et al. Anti-GD2 antibody with GM-CSF, interleukin-2, and isotretinoin for neuroblastoma. N Engl J Med 363, 1324–1334 (2010).

31. Mody R, et al. Irinotecan-temozolomide with temsirolimus or dinutuximab in children with refractory or relapsed neuroblastoma (COG ANBL1221): an open-label, randomised, phase 2 trial. The Lancet Oncology 18, 946–957 (2017).

32. Mody R, et al. Irinotecan, Temozolomide, and Dinutuximab With GM-CSF in Children With Refractory or Relapsed Neuroblastoma: A Report From the Children’s Oncology Group. J Clin Oncol 38, 2160–2169 (2020).

33. van Groningen T, et al. Neuroblastoma is composed of two super-enhancer-associated differentiation states. Nature genetics 49, 1261–1266 (2017).

34. Boeva V, et al. Heterogeneity of neuroblastoma cell identity defined by transcriptional circuitries. Nature genetics 49, 1408–1413 (2017).

35. van Groningen T, et al. A NOTCH feed-forward loop drives reprogramming from adrenergic to mesenchymal state in neuroblastoma. Nature communications 10, 1530 (2019).

36. van Wezel EM, et al. Mesenchymal Neuroblastoma Cells Are Undetected by Current mRNA Marker Panels: The Development of a Specific Neuroblastoma Mesenchymal Minimal Residual Disease Panel. JCO Precision Oncology, 1–11 (2019).

37. Yogev O, et al. In Vivo Modeling of Chemoresistant Neuroblastoma Provides New Insights into Chemorefractory Disease and Metastasis. Cancer research 79, 5382–5393 (2019).

38. van Nes J, Chan A, van Groningen T, van Sluis P, Koster J, Versteeg R. A NOTCH3 transcriptional module induces cell motility in neuroblastoma. Clinical cancer research : an official journal of the American Association for Cancer Research 19, 3485–3494 (2013).

39. Liberzon A, Birger C, Thorvaldsdottir H, Ghandi M, Mesirov JP, Tamayo P. The Molecular Signatures Database (MSigDB) hallmark gene set collection. Cell Syst 1, 417–425 (2015).

40. Ghandi M, et al. Next-generation characterization of the Cancer Cell Line Encyclopedia. Nature 569, 503-508 (2019).

41. Lampson LA, George DL. Interferon-mediated induction of class I MHC products in human neuronal cell lines: analysis of HLA and beta 2-m RNA, and HLA-A and HLA-B proteins and polymorphic specificities. J Interferon Res 6, 257–265 (1986).

42. Reid GS, et al. Interferon-gamma-dependent infiltration of human T cells into neuroblastoma tumors in vivo. Clinical cancer research : an official journal of the American Association for Cancer Research 15, 6602–6608 (2009).

43. Harenza JL, et al. Transcriptomic profiling of 39 commonly-used neuroblastoma cell lines. Sci Data 4, 170033 (2017).

44. Rokita JL, et al. Genomic Profiling of Childhood Tumor Patient-Derived Xenograft Models to Enable Rational Clinical Trial Design. Cell reports 29, 1675–1689 e1679 (2019).

45. Johnson LA, et al. Gene transfer of tumor-reactive TCR confers both high avidity and tumor reactivity to nonreactive peripheral blood mononuclear cells and tumor-infiltrating lymphocytes. J Immunol 177, 6548–6559 (2006).

46. Rajbhandari P, et al. Cross-Cohort Analysis Identifies a TEAD4-MYCN Positive Feedback Loop as the Core Regulatory Element of High-Risk Neuroblastoma. Cancer discovery 8, 582–599 (2018).

47. Valentijn LJ, et al. Functional MYCN signature predicts outcome of neuroblastoma irrespective of MYCN amplification. Proceedings of the National Academy of Sciences of the United States of America 109, 19190–19195 (2012).

48. Henriksen JR, et al. Conditional expression of retrovirally delivered anti-MYCN shRNA as an in vitro model system to study neuronal differentiation in MYCN-amplified neuroblastoma. BMC Dev Biol 11, 1 (2011).

49. Valentijn LJ, Koppen A, van Asperen R, Root HA, Haneveld F, Versteeg R. Inhibition of a new differentiation pathway in neuroblastoma by copy number defects of N-myc, Cdc42, and nm23 genes. Cancer research 65, 3136–3145 (2005).

50. Ushmorov A, Hogarty MD, Liu X, Knauss H, Debatin KM, Beltinger C. N-myc augments death and attenuates protective effects of Bcl-2 in trophically stressed neuroblastoma cells. Oncogene 27, 3424–3434 (2008).

51. Ross RA, Spengler BA, Biedler JL. Coordinate morphological and biochemical interconversion of human neuroblastoma cells. Journal of the National Cancer Institute 71, 741–747 (1983).

52. Dong R, et al. Single-Cell Characterization of Malignant Phenotypes and Developmental Trajectories of Adrenal Neuroblastoma. Cancer Cell 38, 716–733 e716 (2020).

53. Yang X, et al. Induction of caspase 8 by interferon gamma renders some neuroblastoma (NB) cells sensitive to tumor necrosis factor-related apoptosis-inducing ligand (TRAIL) but reveals that a lack of membrane TR1/TR2 also contributes to TRAIL resistance in NB. Cancer research 63, 1122–1129 (2003).

54. Wexler LH, Thiele, C., McClure, L., Chanock, S., Mertins, S., Tsokos, M., Avila, N., Reynolds, J., Ognibene, F., Venzon, D., Castleberry, R., Casper, J., Truitt, R., Yannelli, J., Schwartzentruber, D., Rosenberg, S., Horowitz, M. Adoptive immunotherapy of refractory neuroblastoma with tumor-infiltrating lymphocytes, interferon-gamma, and interleukin-2. In: Proc. Am. Soc. Clin. Oncol. Annu. Meet.) (1992).

55. Molenaar JJ, et al. Sequencing of neuroblastoma identifies chromothripsis and defects in neuritogenesis genes. Nature 483, 589–593 (2012).

56. Koche RP, et al. Extrachromosomal circular DNA drives oncogenic genome remodeling in neuroblastoma. Nature genetics 52, 29–34 (2020).

57. Chuang JH, et al. Preferential involvement of mitochondria in Toll-like receptor 3 agonist-induced neuroblastoma cell apoptosis, but not in inhibition of cell growth. Apoptosis 17, 335–348 (2012).

58. Chuang JH, et al. Differential toll-like receptor 3 (TLR3) expression and apoptotic response to TLR3 agonist in human neuroblastoma cells. J Biomed Sci 18, 65 (2011).

59. Hsu WM, et al. Toll-like receptor 3 expression inhibits cell invasion and migration and predicts a favorable prognosis in neuroblastoma. Cancer Lett 336, 338–346 (2013).

60. Lin LL, Huang CC, Wu CL, Wu MT, Hsu WM, Chuang JH. Downregulation of c-Myc is involved in TLR3-mediated tumor death of neuroblastoma xenografts. Lab Invest 96, 719–730 (2016).

61. Boes M, Meyer-Wentrup F. TLR3 triggering regulates PD-L1 (CD274) expression in human neuroblastoma cells. Cancer Lett 361, 49–56 (2015).

62. Flood BA, Higgs EF, Li S, Luke JJ, Gajewski TF. STING pathway agonism as a cancer therapeutic. Immunol Rev 290, 24–38 (2019).

63. Wang-Bishop L, et al. Potent STING activation stimulates immunogenic cell death to enhance antitumor immunity in neuroblastoma. J Immunother Cancer 8, (2020).

64. Lampkin BC, Levine AS, Levy H, Krivit W, Hammond D. Phase II trial of a complex polyriboinosinic-polyribocytidylic acid with poly-L-lysine and carboxymethyl cellulose in the treatment of children with acute leukemia and neuroblastoma: a report from the Children’s Cancer Study Group. Cancer research 45, 5904–5909 (1985).

65. Hartman LL, et al. Pediatric phase II trials of poly-ICLC in the management of newly diagnosed and recurrent brain tumors. J Pediatr Hematol Oncol 36, 451–457 (2014).

66. Yu AL, et al. Phase I trial of a human-mouse chimeric anti-disialoganglioside monoclonal antibody ch14.18 in patients with refractory neuroblastoma and osteosarcoma. J Clin Oncol 16, 2169–2180 (1998).

67. Weiss WA, Aldape K, Mohapatra G, Feuerstein BG, Bishop JM. Targeted expression of MYCN causes neuroblastoma in transgenic mice. The EMBO journal 16, 2985–2995 (1997).

68. Walton ZE, et al. Acid Suspends the Circadian Clock in Hypoxia through Inhibition of mTOR. Cell 174, 72–87 e32 (2018).

69. Berg S, et al. ilastik: interactive machine learning for (bio)image analysis. Nat Methods 16, 1226–1232 (2019).

70. Kamentsky L, et al. Improved structure, function and compatibility for CellProfiler: modular high-throughput image analysis software. Bioinformatics 27, 1179–1180 (2011).

71. Bolger AM, Lohse M, Usadel B. Trimmomatic: a flexible trimmer for Illumina sequence data. Bioinformatics 30, 2114–2120 (2014).

72. Langmead B, Salzberg SL. Fast gapped-read alignment with Bowtie 2. Nat Methods 9, 357–359 (2012).

73. Li B, Dewey CN. RSEM: accurate transcript quantification from RNA-Seq data with or without a reference genome. BMC Bioinformatics 12, 323 (2011).

74. Satija R, Farrell JA, Gennert D, Schier AF, Regev A. Spatial reconstruction of single-cell gene expression data. Nat Biotechnol 33, 495–502 (2015).

75. McGinnis CS, Murrow LM, Gartner ZJ. DoubletFinder: Doublet Detection in Single-Cell RNA Sequencing Data Using Artificial Nearest Neighbors. Cell Syst 8, 329–337 e324(2019).

76. Tirosh I, et al. Dissecting the multicellular ecosystem of metastatic melanoma by single-cell RNA-seq. Science 352, 189–196 (2016).

77. Bindea G, et al. Spatiotemporal dynamics of intratumoral immune cells reveal the immune landscape in human cancer. Immunity 39, 782–795 (2013).

78. Zhang W, et al. Comparison of RNA-seq and microarray-based models for clinical endpoint prediction. Genome Biol 16, 133 (2015).

