## Supplemental Figures and Tables for "Epigenetic state determines inflammatory sensing in neuroblastoma"

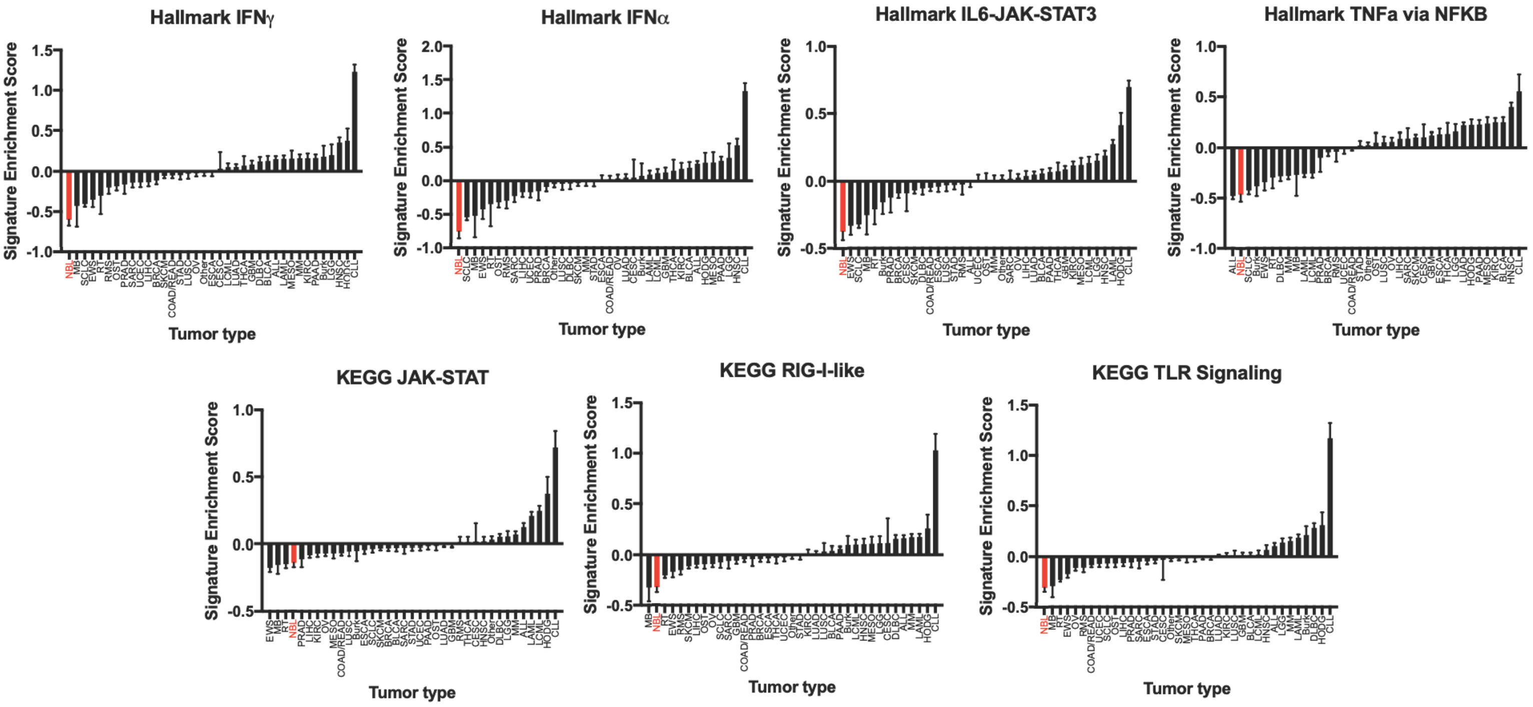


**Figure S1: Enrichment scores of inflammatory signatures in the CCLE**

Enrichment scores across 7 different inflammatory signatures for cell lines in the CCLE, with neuroblastoma highlighted in red. Data downloaded from the Broad CCLE portal (https://portals.broadinstitute.org/ccle). Abbreviations: ALL = acute lymphoblastic leukemia, BRCA = breast invasive carcinoma, Burk = Burkitt lymphoma, CESC = cervical squamous cell carcinoma and endocervical adenocarcinoma, CLL = chronic lymphocytic leukemia, COAD/READ = colon adenocarcinoma/rectum adenocarcinoma, DLBCL = lymphoid neoplasm diffuse large B-cell lymphoma, ESCA = esophageal carcinoma, EWS = Ewing sarcoma, GBM = glioblastoma multiforme, HODG = Hodgkin's lymphoma, HNSC = head and neck squamous cell carcinoma, KIRC = kidney renal clear cell carcinoma, LAML = acute myeloid leukemia, LCML = chronic myelogenous leukemia, LGG = brain lower grade glioma, LIHC = liver hepatocellular carcinoma,

LUAD = lung adenocarcinoma, LUSC = lung squamous cell carcinoma, MB = medulloblastoma, MESO = mesothelioma, MM = multiple myeloma, NBL = neuroblastoma, OST = osteosarcoma, OV = ovarian serous cystadenocarcinoma, PAAD = pancreatic adenocarcinoma, PRAD = prostate adenocarcinoma, RMS = rhabdomyosarcoma, RT = rhabdoid tumor, SARC = sarcoma, SCLC = small cell lung carcinoma, SKCM = skin cutaneous melanoma, STAD = stomach adenocarcinoma, THCA = thyroid carcinoma, UCEC = uterine corpus endometrial carcinoma.

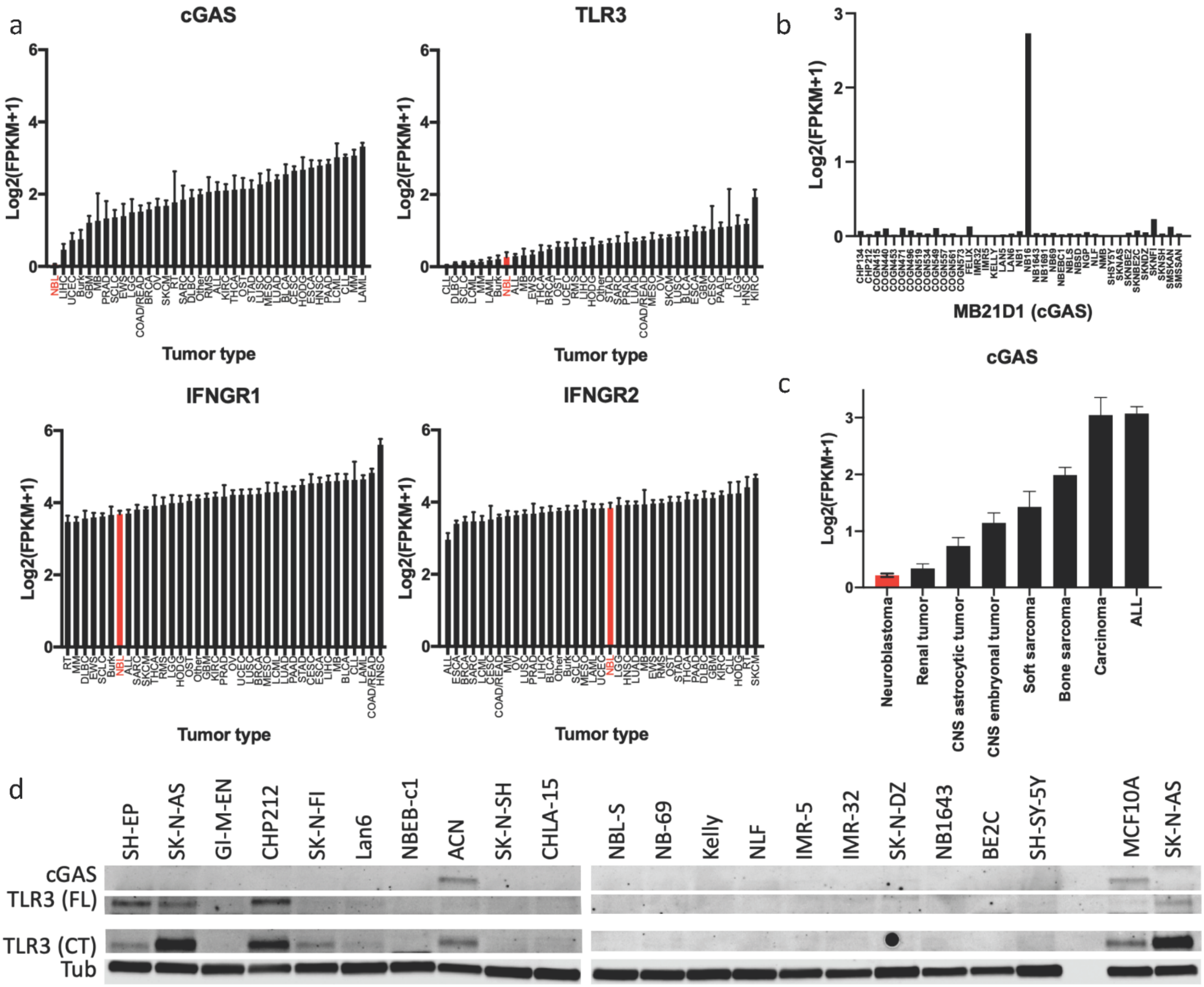


**Figure S2: Expression of inflammatory sensors in the CCLE and in neuroblastoma cell lines**

a) Expression of the indicated genes across tumor types in the CCLE. Data downloaded from the Broad CCLE portal (https://portals.broadinstitute.org/ccle). See Figure S1 for abbreviations. b) Expression of *cGAS* in a neuroblastoma cell line RNA-seq dataset (GSE89413[43]). c) Expression of *cGAS* in a pediatric tumor xenograft dataset, separated by tumor type as indicated. Data from[44], downloaded from PedcBioPortal (https://pedcbioportal.kidsfirstdrc.org). Western blot demonstrating expression of cGAS and TLR3 in the 20 neuroblastoma cell lines used in the current study. Both the full length (FL) and an active C-terminal fragment (CT) of TLR3 are shown. MCF10A cells are shown as a positive control for cGAS.

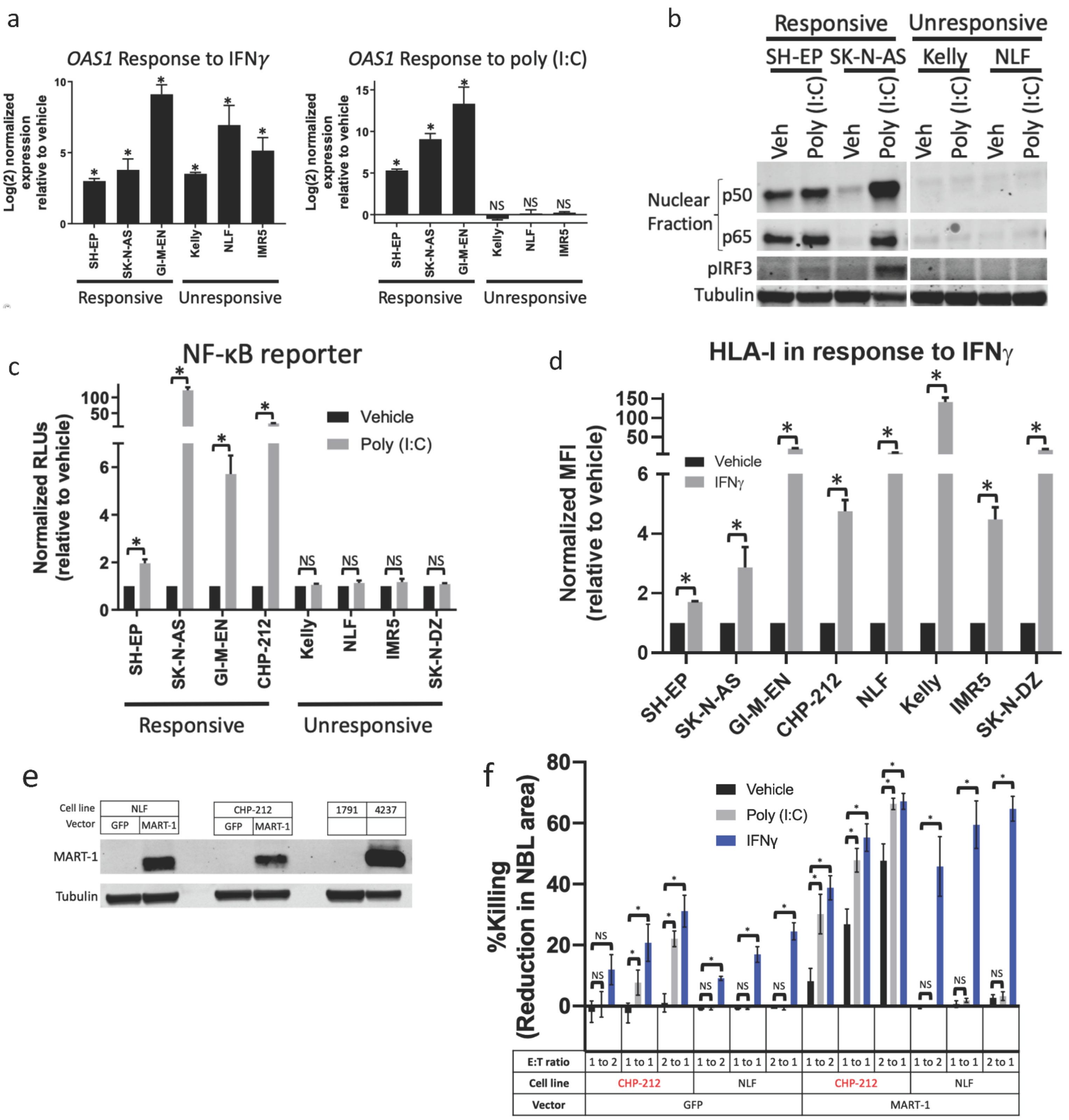


**Figure S3: Additional metrics of differential response to a TLR3 agonist in neuroblastoma cell lines**

a) Change in expression of *OAS1* as measured by qPCR in the indicated neuroblastoma cell lines after treatment with 20ng/mL IFNγ for 24 hours (left) or 30μg/mL of poly (I:C) (right) compared to vehicle control. b) Change in the phosphorylation of IRF3 and the nuclear localization of NF-κB subunits p50 and p65 after treatment of the indicated cell lines with vehicle control or 30μg/mL poly (I:C) for 24 hours. c) Luminescence normalized to vehicle only control in the indicated cell lines transfected with an NF-κB reporter after treatment with vehicle control or 30μg/mL poly (I:C) for 24 hours. d) Surface expression of HLA-I, measured by flow cytometry, in the indicated neuroblastoma cell lines after treatment with vehicle control or 20ng/mL IFNγ for 24 hours. e) Western blot showing expression of MART-1 in NLF and CHP-212 cells after expression of GFP control or MART-1. MART-1 negative (1791) and MART-1 positive (4237) melanoma cells are shown as a positive control. Western blots are representative of results from at least three separate experiments. f) Change in killing of neuroblastoma cells exogenously expressing either GFP control or MART-1 after culture with MART-1 transgenic tTCR-transfected T-cells. Neuroblastoma cells were treated with the indicated agonists for 24 hours then washed and cultured with the T-cells. E:T ratio is the effector (T-cell) to target (neuroblastoma) ratio. Killing was calculated by microscopy-based detection of change in cell area. Two-tailed paired T-test between biological replicates, $^*p<0.05$.

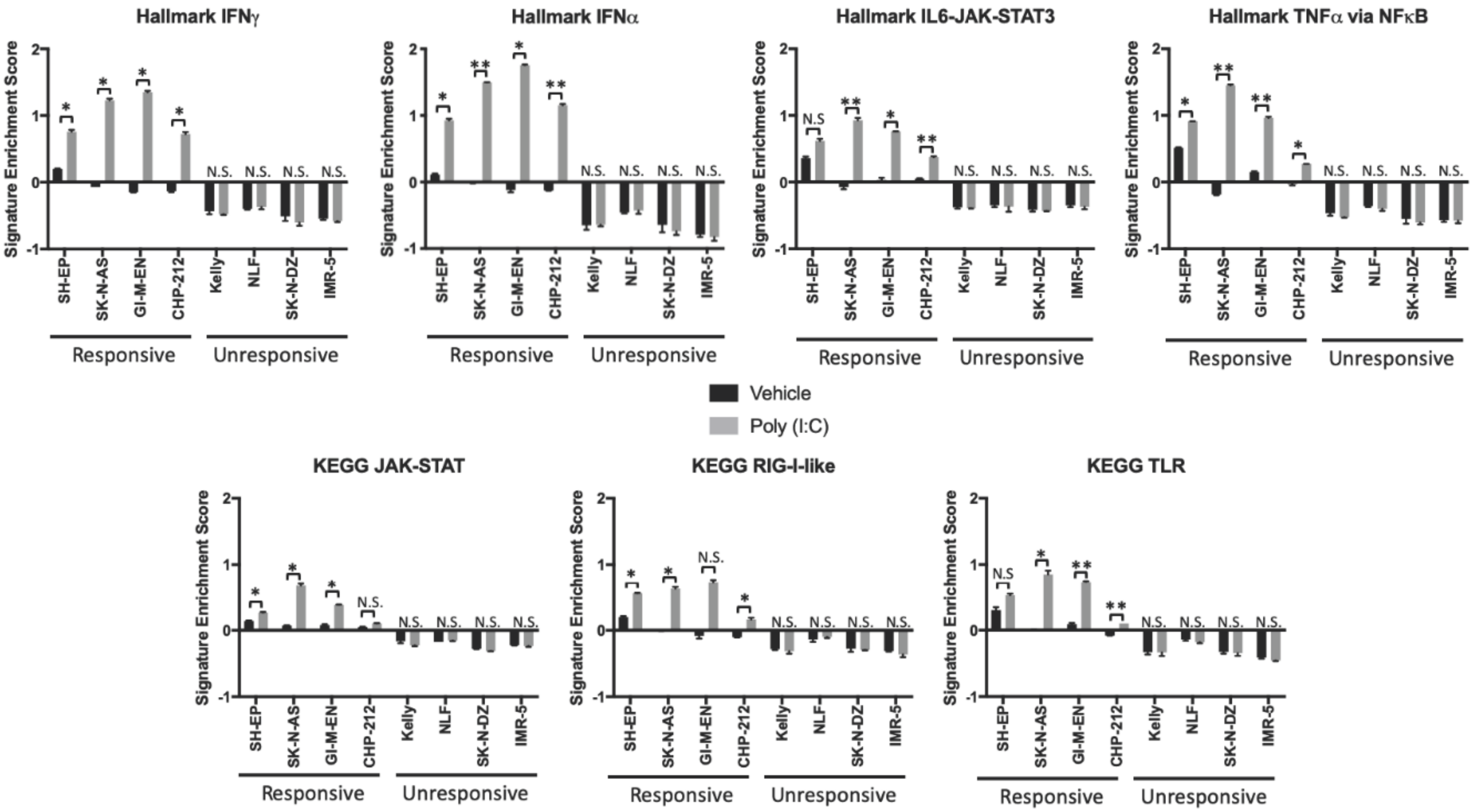


**Figure S4: Changes in gene expression signatures in neuroblastoma cell lines after treatment with a TLR3 agonist**

Relative enrichment of 7 different inflammatory signaling signatures in the indicated cell lines treated with vehicle or 30μg/mL of poly (I:C) for 24 hours as measured by Quantseq. Two-tailed paired T-test between biological replicates showing an increase in the signature upon treatment, *p<0.05, **p<0.01.

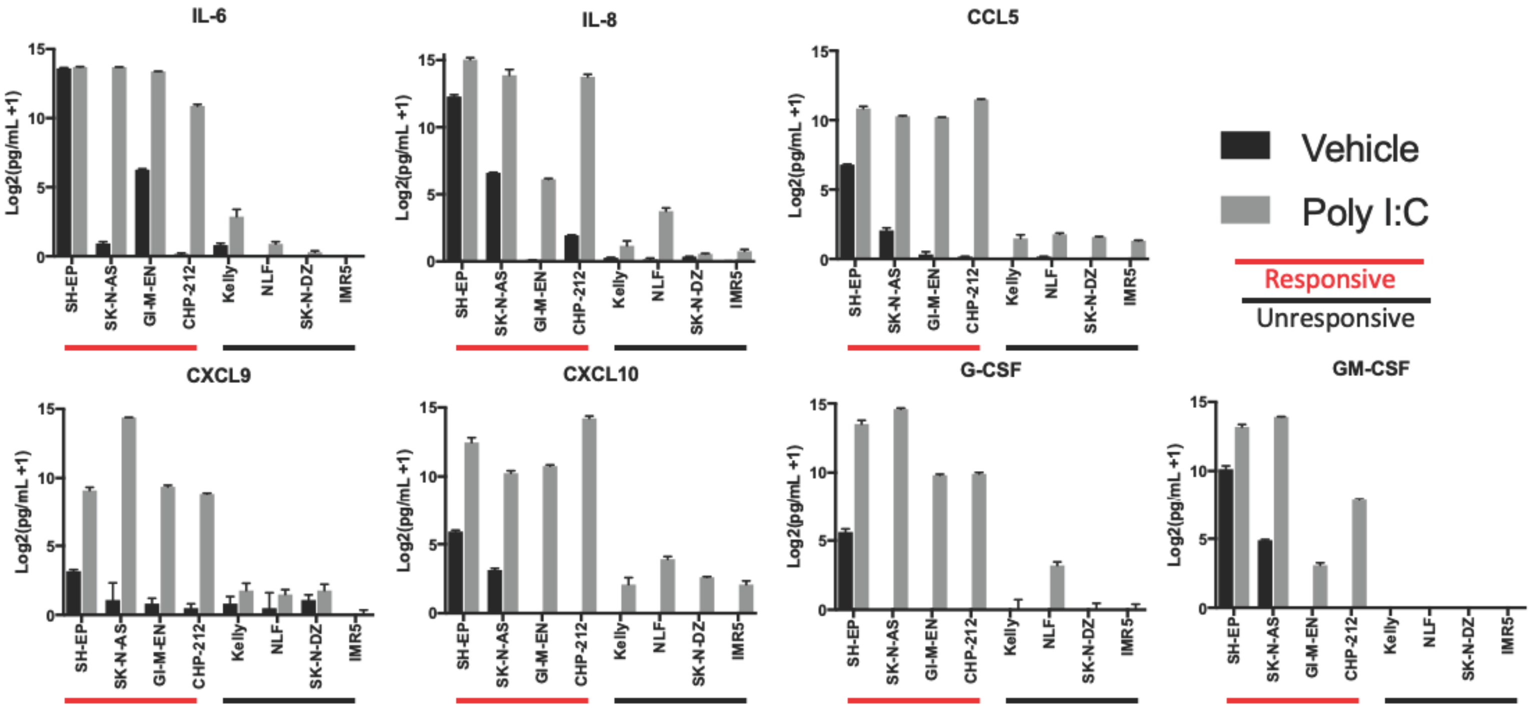


**Figure S5: Changes in cytokine secretion upon treatment with a TLR3 agonist**

Change in cytokines in supernatant of the indicated cell lines after treatment with vehicle or 30μg/mL of poly (I:C) for 24 hours, measured as Log2(pg/mL+1). Comparison between the level of each responsive cell line after treatment is significantly different than that of each unresponsive line ($p<0.02$) for all cytokines shown.

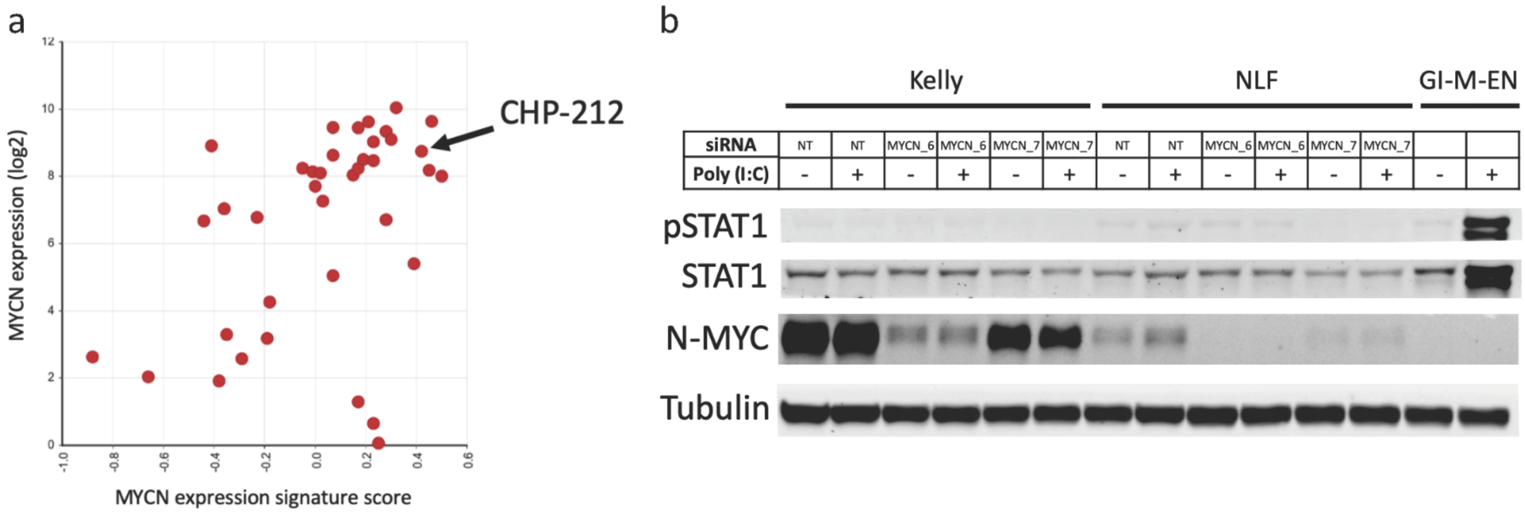


**Figure S6: *MYCN* siRNA does not change TLR3 responsiveness**

a) Comparison of *MYCN* expression and relative enrichment score of a functional MYCN signature[47]. Data from (GSE89413[43]), obtained from and analyzed in R2 (http://r2.amc.nl). b) Western blot showing changes in pSTAT1, STAT1, and MYCN when Kelly or NLF cells were treated with either a control siRNA or one of two different siRNAs targeting *MYCN* for 72 hours, at which point they were then treated with either vehicle or 30μg/mL of poly (I:C) for 24 hours.

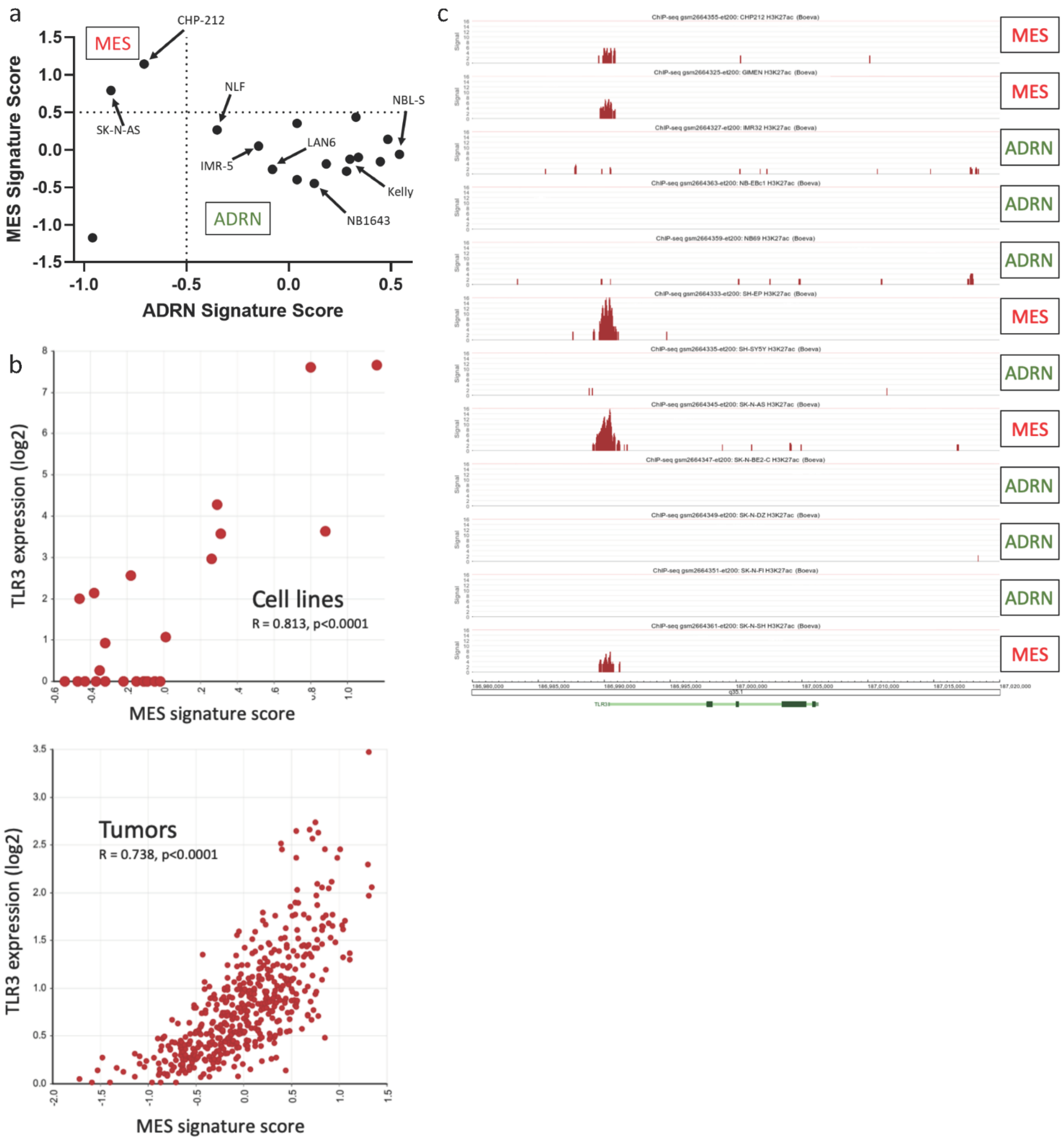


**Figure S7: Classification of neuroblastoma cell lines and relationship between TLR3 and MES expression signature**

a) Comparison of MES and ADRN gene expression signature scores for cell lines used in the current study, data from (GSE89413[43]), signatures from[33]. Dotted lines indicate classification cutoffs used. b) Top - comparison of *TLR3* expression and the relative enrichment score of the MES signature in 23 neuroblastoma cell lines. Data from GSE28019, obtained from and analyzed in R2 (http://r2.amc.nl). Bottom - comparison of *TLR3* expression and the relative enrichment score of the MES signature in 498 neuroblastoma tumors. Data from[78], obtained from and analyzed in R2 (http://r2.amc.nl). c) ChIP-seq for H3K27Ac in the 12 indicated neuroblastoma cell lines surrounding the TLR3 locus. Data from[34], obtained from and analyzed in R2 (http://r2.amc.nl). Y-axis represents the number of reads per 20 million mapped reads.

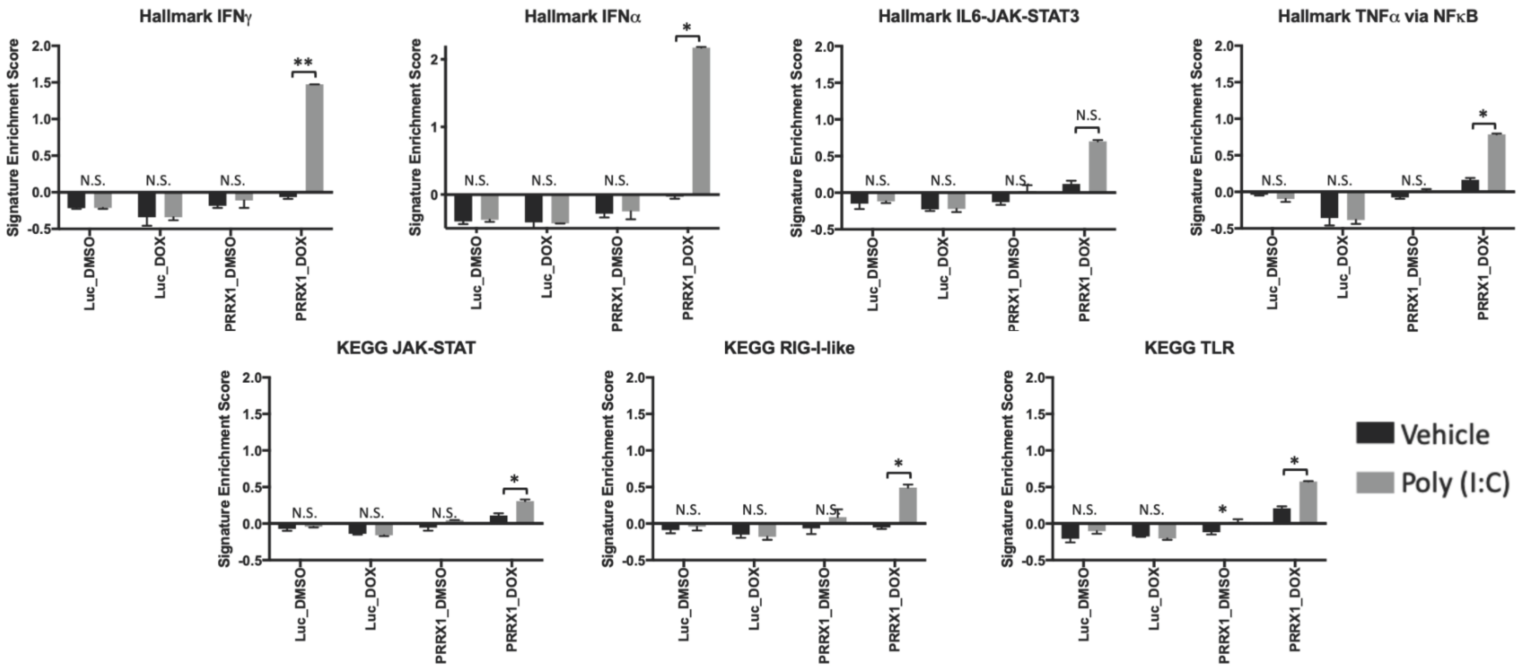


**Figure S8: Effect of PRRX1 expression on the change in gene expression signatures after treatment with a TLR3 agonist**

Relative enrichment of 7 different inflammatory signaling signatures in BE2(c) cells expressing inducible Luciferase control (Luc) or PRRX1 treated with vehicle or doxycycline for 14 days, then treated with vehicle or 30μg/mL of poly (I:C) for 24 hours as measured by Quantseq. Two-tailed paired T-test between biological replicates showing an increase in the signature upon treatment, $*p<0.05$, $**p<0.01$.

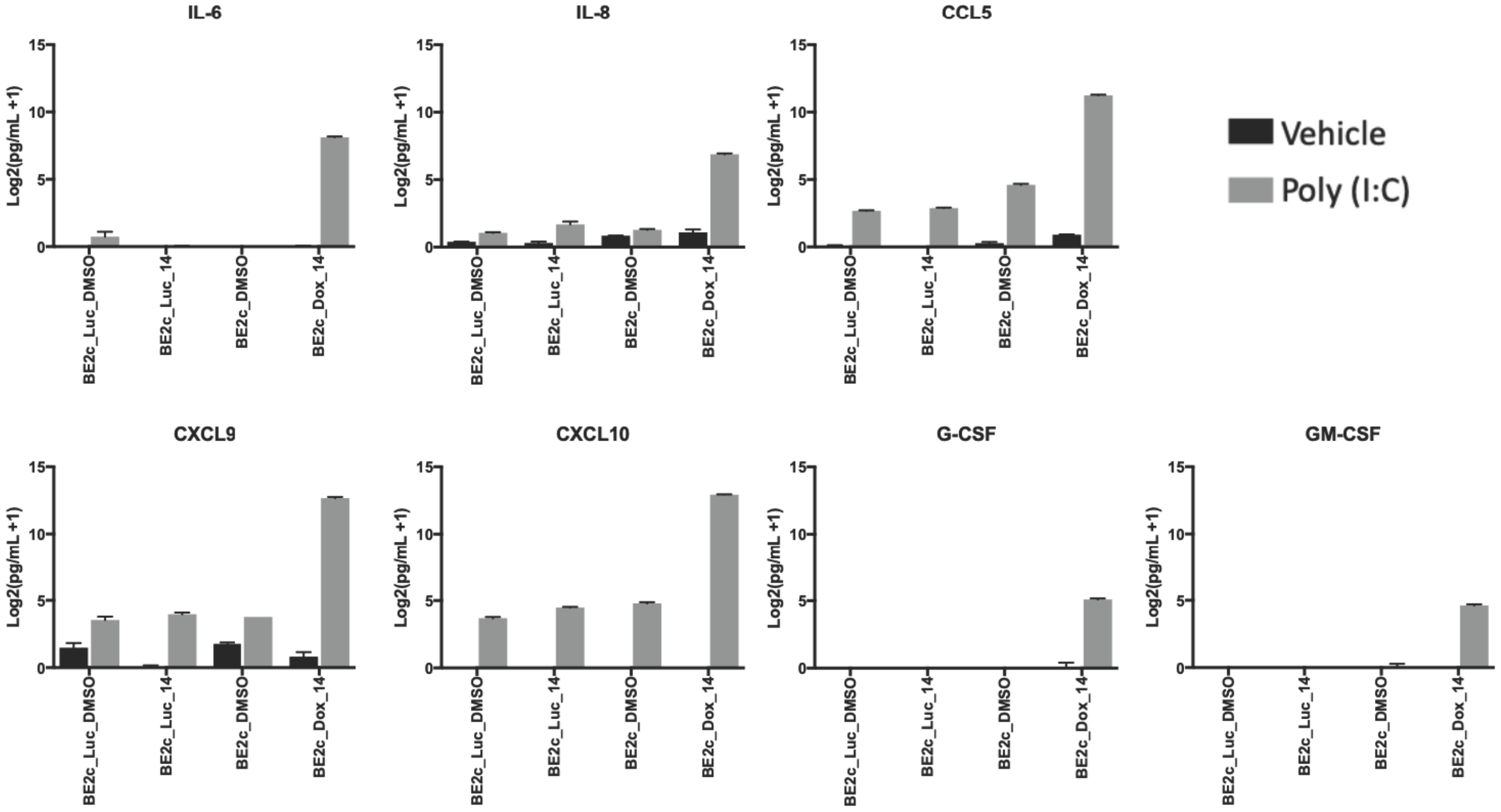


**Figure S9: Changes in cytokine secretion upon treatment with a TLR3 agonist with PRRX1 expression**

Change in cytokines in supernatant of BE2(c) cells expressing inducible Luc or PRRX1 treated with vehicle or dox for 14 days, then treated with vehicle or 30μg/mL of poly (I:C) for 24 hours measured as Log2(pg/mL+1). Comparison between the level of PRRX1 expressing cells treated with dox and poly I:C is significantly different ($p<0.01$) from all other samples ($p<0.01$) for each cytokine shown.

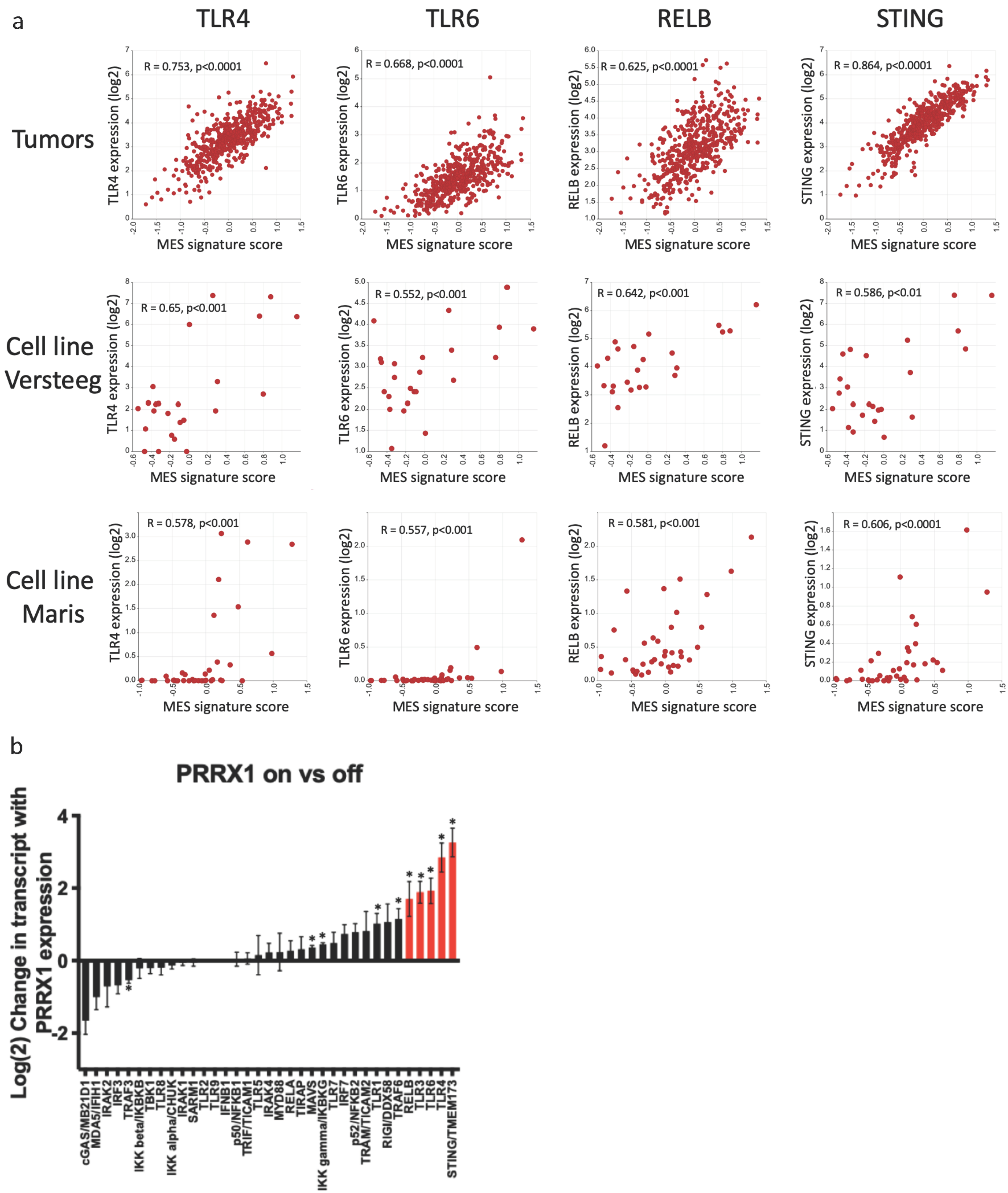


**Figure S10: Relationship between MES signature and additional inflammatory sensors**

a) Comparison of *TLR4, TLR6, RELB, and STING* expression and the relative enrichment score of the MES signature[33]. Top row shows relationship in 498 neuroblastoma tumors (data from[78]). Middle row shows relationship in 23 neuroblastoma cell lines (data from GSE28019). Bottom row shows relationship in 39 neuroblastoma cell lines (data from[43]). All data obtained from and analyzed in R2 (http://r2.amc.nl). b) Change in expression of receptors, adaptors/signaling proteins, and effector transcription factors involved in TLR and other pattern recognition receptor signaling with PRRX1 expression in BE2(c) cells. Cells induced to express PRRX1 were compared to three pooled control conditions (luciferase control vector on/off, PRRX1 vector off). Data from Quantseq analysis. Transcripts highlighted in red were significantly correlated with the MES signature in tumors and in two cell line datasets (shown in panel (a)). Two-tailed T-test *$p<0.05$.

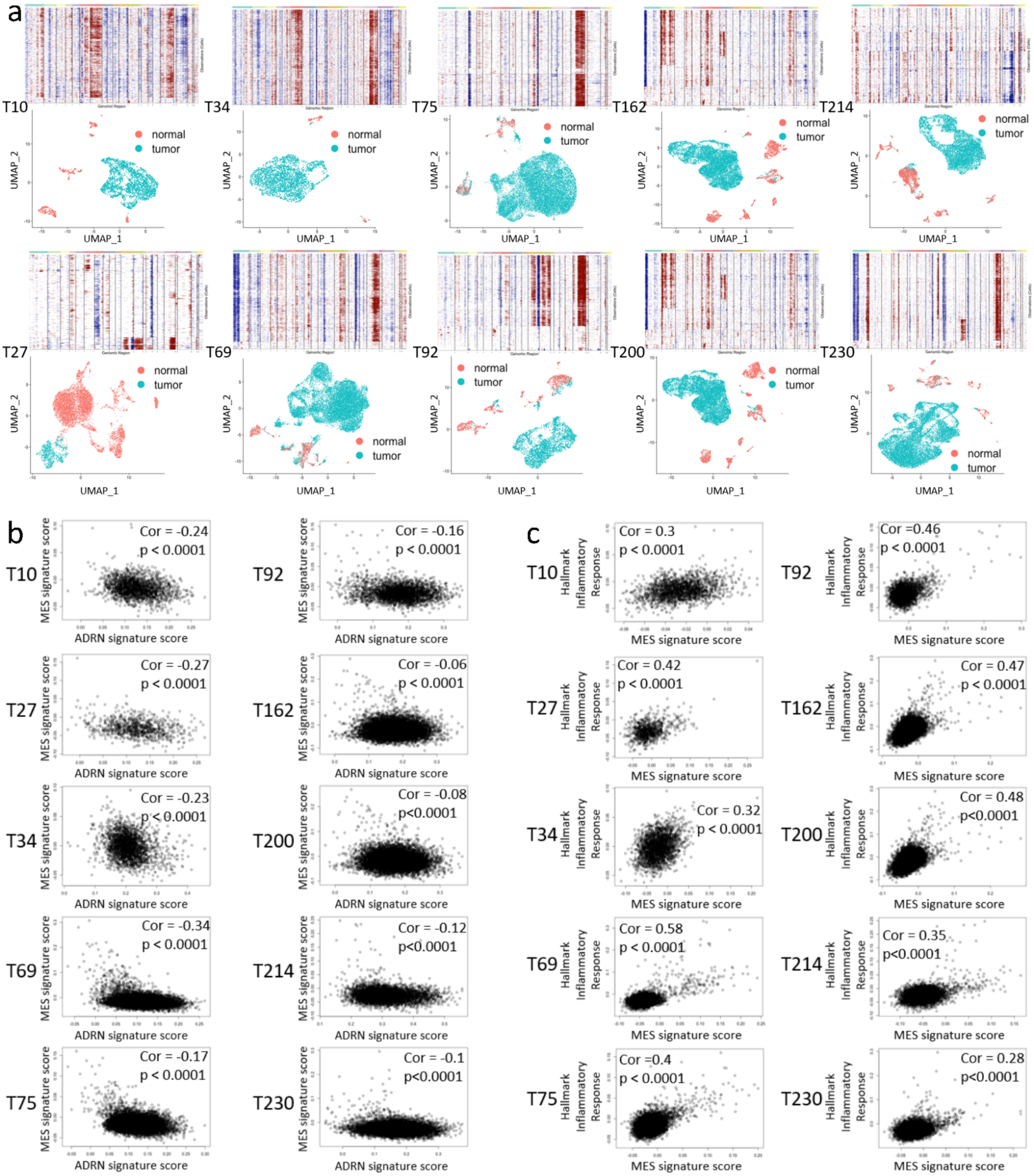


**Figure S11: scRNAseq analysis and data from individual tumors**

a) For each sample, the top plot shows inferred CNV across for each cell (Y-axis) across chromosome location (X-axis). The bottom shows a UMAP plot for each sample with tumors cells assigned based on the CNV analysis in the above plot. Each sample is labelled to the left of the plots. b) Correlation between the MES and ADRN gene signatures[33] in the cells determined to be tumor based on CNV for each tumor sample. c) Correlation between the Hallmark Inflammatory Response signature[39] and the MES signature in the cells determined to be tumor based on CNV for each tumor sample.

**Table S1: Correlations between Mesenchymal gene signatures and pattern recognition signaling gene expression**

| | Versteeg dataset | | Maris dataset | | Tumor RNA-seq | |
|---|---|---|---|---|---|---|
| **Gene name** | **R-value** | **P-value** | **R-value** | **P-value** | **R-value** | **P-value** |
| TLR1 | 0.175 | 0.423 | 0.583 | 0.000122 | 0.696 | 2.06E-73 |
| TLR2 | 0.046 | 0.836 | 0.505 | 0.00123 | 0.741 | 8.89E-88 |
| TLR3 | 0.842 | 4.85E-07 | 0.61 | 0.0000478 | 0.738 | 8.56E-87 |
| TLR4 | 0.65 | 0.000796 | 0.578 | 0.000143 | 0.753 | 3.22E-92 |
| TLR5 | 0.158 | 0.472 | -0.024 | 0.886 | 0.713 | 1.64E-78 |
| TLR6 | 0.552 | 0.00635 | 0.557 | 0.000277 | 0.668 | 1.34E-65 |
| TLR7 | 0.061 | 0.783 | 0.247 | 0.136 | 0.688 | 3.62E-71 |
| TLR8 | 0.297 | 0.169 | Not expressed | Not expressed | 0.603 | 1.24E-50 |
| TLR9 | -0.028 | 0.901 | -0.254 | 0.124 | 0.271 | 7.59E-10 |
| MYD88 | 0.379 | 0.075 | 0.466 | 0.0032 | 0.458 | 3.67E-27 |
| TBK1 | 0.175 | 0.424 | 0.752 | 5.31E-08 | 0.325 | 1.07E-13 |
| TRIF/TICAM1 | 0.462 | 0.026 | 0.621 | 0.0000318 | 0.344 | 2.95E-15 |
| IRAK1 | 0.427 | 0.042 | 0.306 | 0.062 | 0.028 | 0.53 |
| IRAK2 | 0.629 | 0.00129 | 0.365 | 0.024 | 0.627 | 9.01E-56 |
| IRAK4 | 0.609 | 0.00203 | 0.664 | 0.00000555 | 0.707 | 1.3E-76 |
| TRAF3 | 0.165 | 0.453 | 0.332 | 0.041 | 0.233 | 1.51E-07 |
| TRAF6 | -0.495 | 0.016 | 0.153 | 0.359 | -0.105 | 0.019 |
| TRAM/TICAM2 | 0.683 | 0.000327 | 0.147 | 0.378 | 0.683 | 9.29E-70 |
| SARM1 | -0.711 | 0.000141 | -0.059 | 0.726 | -0.039 | 0.387 |
| TIRAP | 0.21 | 0.336 | 0.519 | 0.000849 | 0.123 | 0.0061 |
| IKK alpha/CHUK | -0.067 | 0.76 | 0.673 | 0.00000368 | 0.109 | 0.015 |
| IKK beta/IKBKB | 0.364 | 0.088 | 0.736 | 1.43E-07 | 0.57 | 2.54E-44 |
| IKK gamma/IKBKG | 0.614 | 0.00183 | 0.764 | 2.38E-08 | 0.265 | 1.88E-09 |
| IRF3 | 0.558 | 0.00568 | 0.727 | 0.00000024 | 0.599 | 9.42E-50 |
| IRF7 | 0.119 | 0.587 | 0.359 | 0.027 | 0.451 | 2.51E-26 |
| RELA | 0.353 | 0.099 | 0.696 | 0.0000012 | 0.154 | 0.000577 |
| RELB | 0.642 | 0.000969 | 0.581 | 0.00013 | 0.625 | 3.21E-55 |
| p50/NFKB1 | 5.98 | 0.00059 | 0.785 | 5.24E-09 | 0.673 | 6.67E-67 |
| p52/NFKB2 | 0.484 | 0.019 | 0.77 | 1.6E-08 | 0.713 | 1.76E-78 |
| IFNB1 | 0.015 | 0.946 | 0.497 | 0.00149 | 0.07 | 0.121 |
| RIGI/DDX58 | 0.228 | 0.294 | 0.643 | 0.0000135 | 0.645 | 6.99E-60 |
| MDA5/IFIH1 | 0.871 | 6.26E-08 | 0.551 | 0.000314 | 0.701 | 8.2E-75 |
| cGAS/MB21D1 | -0.148 | 0.5 | 0.018 | 0.915 | 0.73 | 3.92E-84 |
| STING/TMEM173 | 0.586 | 0.0033 | 0.606 | 0.000055 | 0.864 | 5.94E-150 |
| MAVS | 0.182 | 0.405 | 0.57 | 0.000187 | 0.224 | 4.47E-07 |

**Table S2: Cell line sources and culture conditions**

| Cell line | Source | Culture Media |
|---|---|---|
| SH-EP | Michael Hogarty | DMEM, 10%FBS, 1%PS |
| SK-N-AS | Michael Hogarty | DMEM, 10%FBS, 1%PS |
| GI-M-EN | German Collection of Microorganisms and Cell Cultures (DSMZ) | DMEM, 10%FBS, 1%PS |
| CHP-212 | John Maris | EMEM/F12 (1:1 mix), 10%FBS, 1%PS, 2mM L-Glutamine |
| SK-N-FI | John Maris | RPMI, 10% FBS, 1%PS, 2mM L-Glutamine |
| SK-N-SH | Michael Hogarty | RPMI, 10% FBS, 1%PS, 2mM L-Glutamine |
| ACN | Interlab Cell Line Collection (ICLC) | RPMI, 10% FBS, 1%PS, 2mM L-Glutamine, 1mM sodium pyruvate |
| NBEBc1 | Michael Hogarty | RPMI, 10% FBS, 1%PS, 2mM L-Glutamine |
| LAN6 | Michael Hogarty | RPMI, 10% FBS, 1%PS, 2mM L-Glutamine |
| NBL-S | John Maris | RPMI, 10% FBS, 1%PS, 2mM L-Glutamine |
| NB69 | John Maris | RPMI, 10% FBS, 1%PS, 2mM L-Glutamine |
| Kelly | Michael Hogarty | DMEM, 10%FBS, 1%PS |
| NLF | Michael Hogarty | DMEM, 10%FBS, 1%PS |
| IMR-5 | Michael Hogarty | DMEM, 10%FBS, 1%PS |
| IMR-32 | Michael Hogarty | RPMI, 10% FBS, 1%PS, 2mM L-Glutamine |
| SK-N-DZ | John Maris | RPMI, 10% FBS, 1%PS, 2mM L-Glutamine |
| NB1643 | Michael Hogarty | RPMI, 10% FBS, 1%PS, 2mM L-Glutamine |
| BE2(c) | Michael Hogarty | RPMI, 10% FBS, 1%PS, 2mM L-Glutamine |
| SH-SY-5Y | Michael Milone | DMEM, 10%FBS, 1%PS |
| CHLA-15 | Michael Hogarty | IMDM, 20% FBS, 1% PS, 2mM L-Glutamine, 0.1% ITS premix (Insulin, Transferrin, Selenious Acid) |
| SH-EP MYCN-ER | Michael Hogarty | DMEM, 10%FBS, 1%PS |
| SK-N-AS MYCN-ER | Linda Valentijn | DMEM, 10%FBS, 1%PS |
| BE2c inducible shRNA | Marie Arsenian-Henriksson | EMEM/F12 (1:1 mix), 10%FBS, 1%PS, 1% Non-essential amino acids, 2mM L-Glutamine |
